# Type 1 piliated uropathogenic *Escherichia coli* hijack the host immune response by binding to CD14

**DOI:** 10.1101/2021.10.18.464770

**Authors:** Kathrin Tomasek, Alexander Leithner, Ivana Glatzova, Michael S. Lukesch, Călin C. Guet, Michael Sixt

## Abstract

A key attribute of persistent or recurring bacterial infections is the ability of the pathogen to evade the host’s immune response. Many *Enterobacteriaceae* express type 1 pili, a pre-adapted virulence trait, to invade host epithelial cells and establish persistent infections. However, the molecular mechanisms and strategies by which bacteria actively circumvent the immune response of the host remain poorly understood. Here, we identified CD14, the major co-receptor for lipopolysaccharide detection, on dendritic cells as a previously undescribed binding partner of FimH, the protein located at the tip of the type 1 pilus of *Escherichia coli*. The FimH amino acids involved in CD14 binding are highly conserved across pathogenic and non-pathogenic strains. Binding of pathogenic bacteria to CD14 lead to reduced dendritic cell migration and blunted expression of co-stimulatory molecules, both rate-limiting factors of T cell activation. While defining an active molecular mechanism of immune evasion by pathogens, the interaction between FimH and CD14 represents a potential target to interfere with persistent and recurrent infections, such as urinary tract infections or Crohn’s disease.

## Introduction

Cells of metazoan organisms constantly interact with a wide diversity of bacterial species that populate the host organism internally as well as externally. Therefore the host immune response is faced with a non-trivial problem – accommodate beneficial commensals and remove harmful pathogens (Hooper & Macpherson, 2010). The difficulty of this task lies in the fact that most of the molecular patterns, such as lipopolysaccharide (LPS) or surface organelles such as pili and flagella, are conserved between commensal and pathogenic bacteria. The main difference between commensal and pathogenic strains is the ability of the latter to hijack host cell functions for their own benefit (Magalhaes *et al,* 2007). Consequently, the host immune system uses complex discrimination strategies, such as spatial compartmentalization of the receptors recognizing pathogen signatures and concurrent sensing of molecular patterns associated with host-damage. The host discrimination ability is not always perfect, as pathogens can persist asymptomatically in the host for long periods of time or cause symptomatic acute or chronic infections in some individuals, but not in others (Grant & Hung, 2013). For example, the commensal bacterium *Escherichia coli*, one of the main residents of the mammalian intestine, occasionally causes clinical infections, especially when the bacteria acquire virulence traits and thus manage to populate extraintestinal host niches (Magalhaes *et al,* 2007; Leimbach *et al,* 2013). Usually disease progression does not pose a dead end to such opportunistic pathogens, since bacteria are shed in high numbers during the infection, allowing the pathogen to populate new environments, a new host or even recolonize their original host niche – the intestine (Donnenberg, 2013).

In principle, any commensal *E. coli* has the potential to evolve into a pathogen by acquiring virulence traits (Wirth *et al,* 2006). Those traits can be either adapted, namely evolved specifically to increase the fitness of a pathogen during the infection, or pre-adapted, meaning even though they increase the fitness of the pathogen they were originally evolved for a non-virulent function (Donnenberg, 2013). Examples of adapted traits are the acquisition of pathogenicity islands through horizontal gene transfer (Oelschlaeger et al., 2002) or pathoadaptive mutations, such as changes in the LPS, flagellum or pili components (Weissman et al., 2006; Donnenberg, 2013). A prime example of a pre-adapted trait are type 1 pili. Both commensal and pathogenic *E. coli* (Shawki & McCole, 2017; Croxen & Finlay, 2010), but also other *Enterobacteriaceae* (Struve *et al,* 2008; Kolenda *et al,* 2019), use type 1 pili to adhere to host cells in their respective ecological niches (Spaulding *et al,* 2017). Type 1 pili undergo a constant and reversible change in their expression due to phase variation which allows populations of isogenic bacteria to exhibit controlled genotypic and phenotypic variation (Bayliss, 2009). In the case of type 1 pili, site-specific recombinases place the *fimA* promoter either in the phase-ON or phase-OFF orientation, resulting in piliated or non-piliated bacteria (Schilling *et al,* 2001). UPECs have mastered the use of phase variation as a remarkable genetic mechanism of plasticity for their advantage, since type 1 pili expression is tightly regulated by the host environment. For example, growing UPECs *in vitro* in human urine locks expression in the phase-OFF state (Greene *et al,* 2015), whereas adhesion to host cells has been shown to lock expression in the phase-ON state (Greene *et al,* 2015; Lim *et al,* 1998). Phase variation greatly adds to the virulence and fitness of UPECs by generating heterogeneity among the bacterial population where individual cells switch back and forth between the type 1 piliated and non-piliated phenotype (Bayliss, 2009; Wright *et al,* 2007). Type 1 pili are necessary for the persistent and therefore recurring infection of the bladder (Hunstad & Justice, 2010; Wright *et al,* 2007), but also other persistent bacterial infections (Shawki & McCole, 2017).

Several mechanisms are known to contribute to persistent or recurring infections, one of which is the residing of pathogenic bacteria within a protected niche (Grant & Hung, 2013). Such niches can be physical structures when the pathogen invades host cells to hide from the immune response (Grant & Hung, 2013; Donnenberg, 2013). For example, UPECs invade host epithelium cells in the bladder using type 1 pili, propagate intracellularly and compromise the host defense barriers (Hunstad & Justice, 2010; Martinez *et al,* 2000). Strikingly, also the host immune response may unintentionally create protected niches. For example, bladder residing macrophages sequester UPECs before antigen-presenting cells, such as dendritic cells (DCs), come into action (Mora-Bau *et al,* 2015).

DCs are the key cell type connecting the innate and adaptive immune response (Mellman & Steinman, 2001). Distinct subtypes of DCs, expressing different surface receptors (Merad et al., 2013), reside in every tissue of the host and DCs were also identified at the epithelial junction of the bladder (Schilling et al., 2003). In response to an infection, these resident DCs together with newly recruited inflammatory DCs sense and ingest pathogens. Subsequently, they start to secrete immune modulatory cytokines and migrate from the site of infection to the draining lymph nodes where they present acquired and processed antigens to T cells, thus triggering the adaptive immune response. However, even if the host immune system manages to detect the pathogen, eradication of the infection is not guaranteed. Pathogens utilize several strategies to subvert innate and adaptive immunity, thereby avoiding clearance by the host and establishing persistent infections (Donnenberg, 2013). For example, UPECs were shown to interfere at the interface between innate and adaptive immunity: after contact with UPECs during urinary tract infections (UTIs), tissue resident mast cells secrete high amounts of interleukin-10 (IL-10), an immuno-suppressive cytokine (Chan *et al,* 2013), that drives the differentiation of regulatory T cells (Hsu *et al,* 2015). The combined effect of subverting the host immune response and establishing a protected niche inside the host greatly increases the ability of pathogens to cause persistent infections.

Here, we asked whether type 1 pili play a role in manipulating host immunity and thereby facilitate recurring bacterial infections by dissecting the underlying molecular mechanism between the interaction of uropathogenic *E. coli* and dendritic cells.

## Results

### Type 1 piliated UPEC inhibit T cell activation and proliferation by decreasing expression of co-stimulatory molecules on dendritic cells

To study the influence of type 1 pili on the adaptive immune system, we engineered bacteria with and without pili. The *fim* genes that produce the molecular components of the type 1 pilus are part of an operon on the *E. coli* chromosome (Figure 1A and B) whose expression is regulated by phase-variation. We generated stable phase-locked bacterial mutants by locking the fim switch (*fimS*) either in the phase-ON orientation, resulting in constitutive expression of the type 1 pilus, or in the phase-OFF orientation, blocking expression of the pilus (hereafter simply termed ON and OFF, respectively). To achieve this, we deleted the 9 bp long recognition site for the site-specific recombinases FimB and FimE in the internal repeat region upstream of the *fimS* element in either orientation (Figure 1C). The presence and absence of type 1 pili was confirmed by electron microscopy and yeast-agglutination assay (Figure 1D). Additionally, electron microscopy indicates absence of any other pili, such as p fimbriae, under the chosen growth conditions since the OFF mutant appears bald (Figure 1D, right panel).

**Figure 1:**
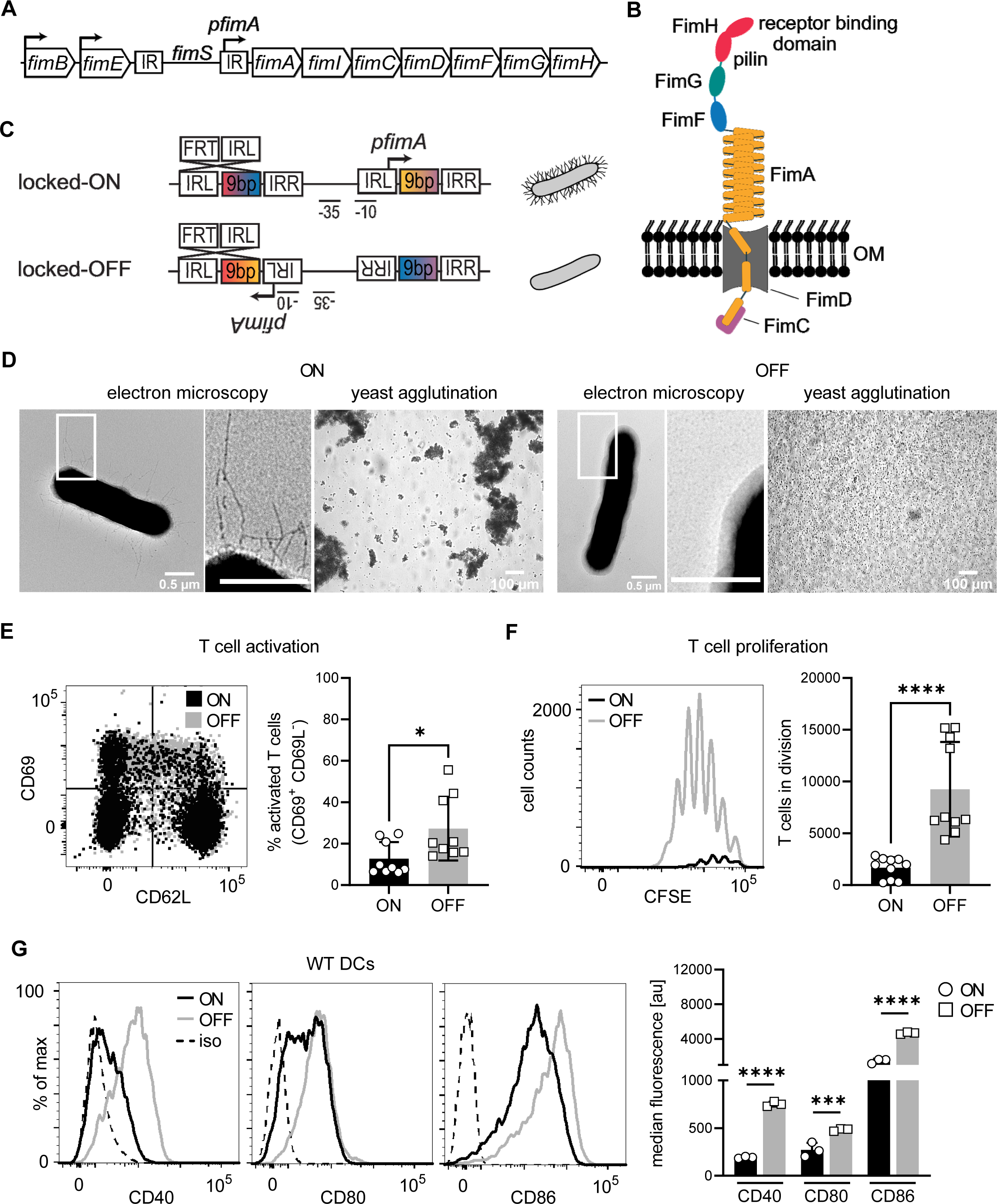
Type 1 piliated UPECs inhibit T cell activation and proliferation by decreasing co-stimulatory molecules on DCs. A. Type 1 pili genes are expressed from the *fim* operon. Phase variation of the *fim* switch (*fimS*) harboring *fimA* promoter (*pfimA*) drives expression. *fimB* and *fimE* genes express site-specific recombinases, FimB and FimE respectively, inverting the *fimS* region by binding to the inverted repeats (IR). B. Type 1 pili consist of several repeating units of the rod protein FimA, two adaptor proteins FimF and FimG, and the tip protein FimH. Fimbrial- and a receptor-binding domain of the two-domain FimH protein are shown. FimD, outer membrane usher. FimC, chaperone. C. Phase-locked mutants were generated by deleting the 9 bp recognition site for the site-specific recombinases in the left inverted repeat region of the *fimS* in either ON or OFF orientation by introducing FRT sites resulting in piliated and non-piliated bacteria (see Methods). D. ON (left panel) and OFF mutants (right panel). Electron microscopy images, with zoomed in regions (white boxes) marked in inlays, and yeast agglutination assay. E. Dot plot of CD69 and CD62L expression on T cells after co-culture with ON (black) and OFF (grey) stimulated DCs (left panel). Quantification of CD69^+^ CD62L^-^ T cell frequencies (right panel) (4 biological replicates). F. CFSE dilution profile of T cells after 96h of co-culture with ON (black) and OFF (grey) stimulated DCs (left). Quantification of T cells in division (right panel) (3 biological replicates). G. Expression level of co-stimulatory molecules (CD40, CD80, CD86) of WT DCs after stimulation with ON (black) and OFF mutants (grey) (left panel; iso – isotype control). Quantification of median fluorescence values of co-stimulatory molecules (right panel) (3 biological replicates). * p<0.1, *** p<0.01, **** p<0.001 by Student’s t test (E, F) and by Holm-Sidak t test (G); data are represented as means ± SD

Since the adaptive immune response seems to be limited during persistent or recurring infections (Magalhaes *et al,* 2007), such as recurring bladder infections (Abraham & Miao, 2015), we asked if activation and proliferation of naïve T cells *in vitro* is altered upon stimulating DCs with the genetically constructed UPEC ON or OFF mutants. The activation of Ovalbumin (OVA) specific CD4^+^ T cells (Barnden et al., 1998), as assessed by CD69 receptor upregulation and CD62L receptor downregulation, was massively reduced when DCs were stimulated with OVA and UPEC ON mutants, compared to OVA and UPEC OFF mutants (Figure 1E). Accordingly, the number of proliferating T cells was also strongly decreased (Figure 1F).

Cellular identity, such as the CD11c surface marker and surface levels of MHCII, the hallmark of DC activation, was only very mildly altered upon exposure to UPEC ON (Supplementary Figure 1C and D). Beyond presentation of MHCII, the expression of co-stimulatory molecules on DCs is decisive for effective T cell priming and differentiation into effector cells (Banchereau & Steinman, 1998). We therefore analyzed surface expression of CD40, CD80 and CD86 after ON and OFF stimulation and found significantly decreased levels after ON stimulation (Figure 1G).

These data suggest that type 1 piliated UPECs prevent effective activation of the adaptive immune response by decreasing expression of co-stimulatory molecules on DCs and thus restrict their ability to activate T cells.

### Type 1 piliated UPEC decrease dendritic cell migratory capacity and increase T cell contact times by triggering integrin activation

Beyond presentation of the antigen in the context of co-stimulatory factors, two additional cell biological parameters are essential for the priming of T cells. First, DCs have to migrate from the site of antigen encounter and uptake (usually peripheral tissues) to the site of antigen presentation (e.g. lymph nodes), where they meet T cells. This migration step is directionally guided by chemokines and highly efficient even in the absence of adhesive interactions with the extracellular matrix (Lämmermann *et al,* 2008). Second, T cells have to physically interact with DCs in order to probe their surface for antigen presentation and co-stimulation. Hence, the contact dynamics between DCs and T cells are essential parameters determining T cell activation and proliferation (Bousso, 2008).

We first investigated DC-T cell contacts using live cell microscopy and found that UPEC ON mutants increased the antigen specific contact times between DCs and naïve CD4^+^ T cells, compared to UPEC OFF mutants. Even after 6 h of co-culture a large fraction of T cells was unable to dissociate from DCs (Figure 2A). This was specific to antigen-bearing DCs, as in the absence of OVA contact times were short and indistinguishable between ON and OFF stimulation. Activation of β2 integrins, specifically CD11b, on DCs was shown to increase the duration of cell-cell contacts by binding to its counter receptor ICAM-1 on T cells, leading to a decrease in the activation of T cells (Varga *et al,* 2007). We therefore investigated if the activation status of CD11b on DCs was affected after stimulation with ON mutants. We analyzed total and active levels of CD11b using the activation-independent antibody M1/70 and CBRM1/5 antibody, which recognizes the active conformation of human, but also mouse CD11b ((Oxvig *et al,* 1999) and (Supplementary Figure 2A)). We found that UPEC ON stimulation shifted CD11b to the active conformation when compared to UPEC OFF or LPS stimulation (Figure 2B and Supplementary Figure 2A).

**Figure 2:**
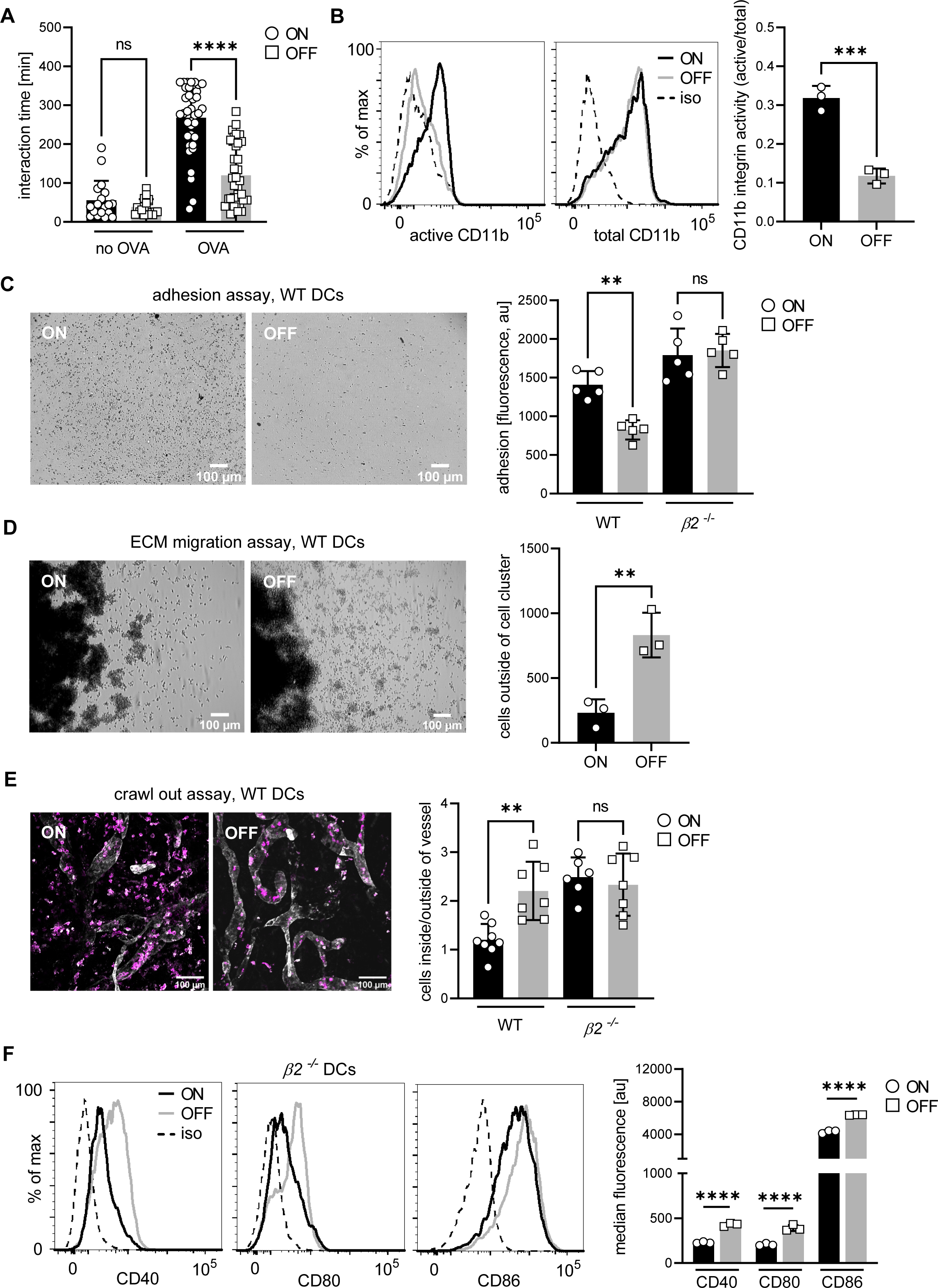
Over-activation of integrins increases DC-T cell contact time and adhesion of DCs to extracellular matrixes leading to decreased migratory capacity. A. Interaction time between ON and OFF stimulated DCs and T cells in the presence and absence of OVA peptide (2 biological replicates). B. Histograms of active and total CD11b integrin after ON (black) and OFF (grey) stimulation of DCs (left panel; iso – isotype control). Quantification of CD11b activity (active/total levels of CD11b) (right panel) (3 biological replicates). (see also Supplementary Figure 2A) C. Adhesion assay of WT DCs after ON and OFF stimulation (left panel). Quantification of fluorescence signals, proxy for adherent cells, after ON (black) and OFF (grey) stimulation of WT and *β2^-/-^* DCs (right panel) (5 biological replicates). (see also Supplementary Figure 2C) D. Extracellular matrix (ECM) migration assay of WT DCs after ON and OFF stimulation (left panel). Quantification of individual cells outside of cell cluster (right panel) (3 biological replicates). E. Ear crawl out assay of WT DCs after ON and OFF stimulation. Endogenous DCs stained with anti-MHCII (magenta). Lymph vessels stained with anti-LYVE-1 (white) (left panel). Quantification of cells inside over outside of lymph vessel after ON (black) and OFF (grey) stimulation of WT and *β2^-/-^* DCs (right panel) (3 biological replicates). (see also Supplementary Figure 2D). F. Expression level of co-stimulatory molecules (CD40, CD80, CD86) of *β2^-/-^* DCs after stimulation with ON (black) and OFF mutants (grey) (left panel; iso – isotype control). Quantification of median fluorescence values of co-stimulatory molecules (right panel) (3 biological replicates). ns, not significant, ** p<0.05, *** p<0.01, **** p<0.001 by one-way ANOVA followed by Dunnett’s multiple comparisons (A, C, E) and by Student’s t test (B, D and F); data are represented as means ± SD

In line with the finding that type 1 piliated UPEC enhanced CD11b activity, ON mutants triggered tight adhesion of DCs to serum-coated surfaces (Figure 2C). This surface immobilization was integrin mediated, as for *β2* integrin knockout DCs the differential adhesion was lost (Figure 2C and Supplementary Figure 2C). (Notably, *β2* integrin deficient DCs showed increased β1 integrin-mediated background-binding (Supplementary Figure 2B)).

We next asked whether UPEC ON stimulation interferes with the migratory capacity of DCs. We performed *in vitro* migration assays where DCs migrate in cell derived matrices (Kaukonen *et al,* 2017) and found diminished migration after stimulation with ON mutants when compared to OFF mutants (Figure 2D). The same effect on the migratory capacity we observed in crawl out assays in physiological tissue (Stösel *et al,* 1997). Here, the ventral halves of explanted mouse ears were repeatedly exposed to either UPEC ON or OFF bacteria during 48 h. After stimulation with ON mutants, fewer endogenous DCs were found inside lymph vessels as compared to OFF stimulation (Figure 2E). In ears harvested from β2 knockout mice the migration defect was rescued and similar levels of *β2*^-/-^ DCs migrated into the lymph vessels after ON and OFF stimulation (Figure 2E and Supplementary Figure 2D).

These data suggest that hyperactive CD11b hinders both migration and T cell activation of DCs by immobilizing them to extracellular matrix proteins like fibrinogen or cellular ligands like ICAM-1. This integrin gain of function phenotype is in line with findings that pharmacological approaches which activate integrins have stronger *in vivo* effects on leukocyte migration than approaches to inhibit integrin function (Maiguel *et al,* 2011). To test if immobilization by hyperactive CD11b is indeed causative for the loss of migration upon UPEC ON stimulation, we measured migration of DCs in 3D collagen gels. Here, DCs efficiently migrate in an integrin-independent manner (Lämmermann *et al,* 2008) and hyperactive CD11b, which does not bind to collagen 1, should not be able to immobilize the cells. We found that the migration speed of ON and OFF stimulated DCs in a 3D collagen assay was indistinguishable (Supplementary Figure 2E).

Finally, we asked if activation of β2 integrins maps upstream of the observed detrimental effect on co-stimulatory molecule expression after stimulation with UPEC ON mutants. *β2*^-/-^ DCs still expressed lowered levels of co-stimulatory molecules after ON stimulation when compared to OFF stimulation (Figure 2F), suggesting that downregulation of co-stimulatory molecules does not depend on integrin activation.

Taken together, type 1 piliated UPEC increase integrin activity on DCs leading to increased cell-cell and cell-matrix attachment. This causes prolonged interaction times with T cells and impaired migratory capacity. Together, with the above finding of reduced co-stimulatory molecule expression, these data demonstrate that type 1 piliated UPEC target three functional hallmarks of DCs that are critical for the activation of naïve T cells – (i) migration of DCs to the lymph node, (ii) the physical interaction with T cells and (iii) the expression of co-stimulatory molecules.

### The GPI-anchored glycoprotein CD14 binds FimH, making it a novel target for type 1 pili

To find the interaction partner on the host cell that leads to integrin activation, we first investigated the role of the major immune cell receptor TLR4 which has been previously suggested as a receptor for the FimH protein of type 1 pili (Mossman *et al,* 2008). We analyzed the adhesion and migration behavior of *tlr4^-/-^* DCs after stimulation with UPEC ON mutants and found that both behaviors were still affected (Figure 3A-B and Supplementary Figure 3A). Our results, together with recent findings in *Salmonella* (Uchiya *et al,* 2019), suggest that other receptors besides TLR4 could serve as molecular targets of type 1 pili.

**Figure 3:**
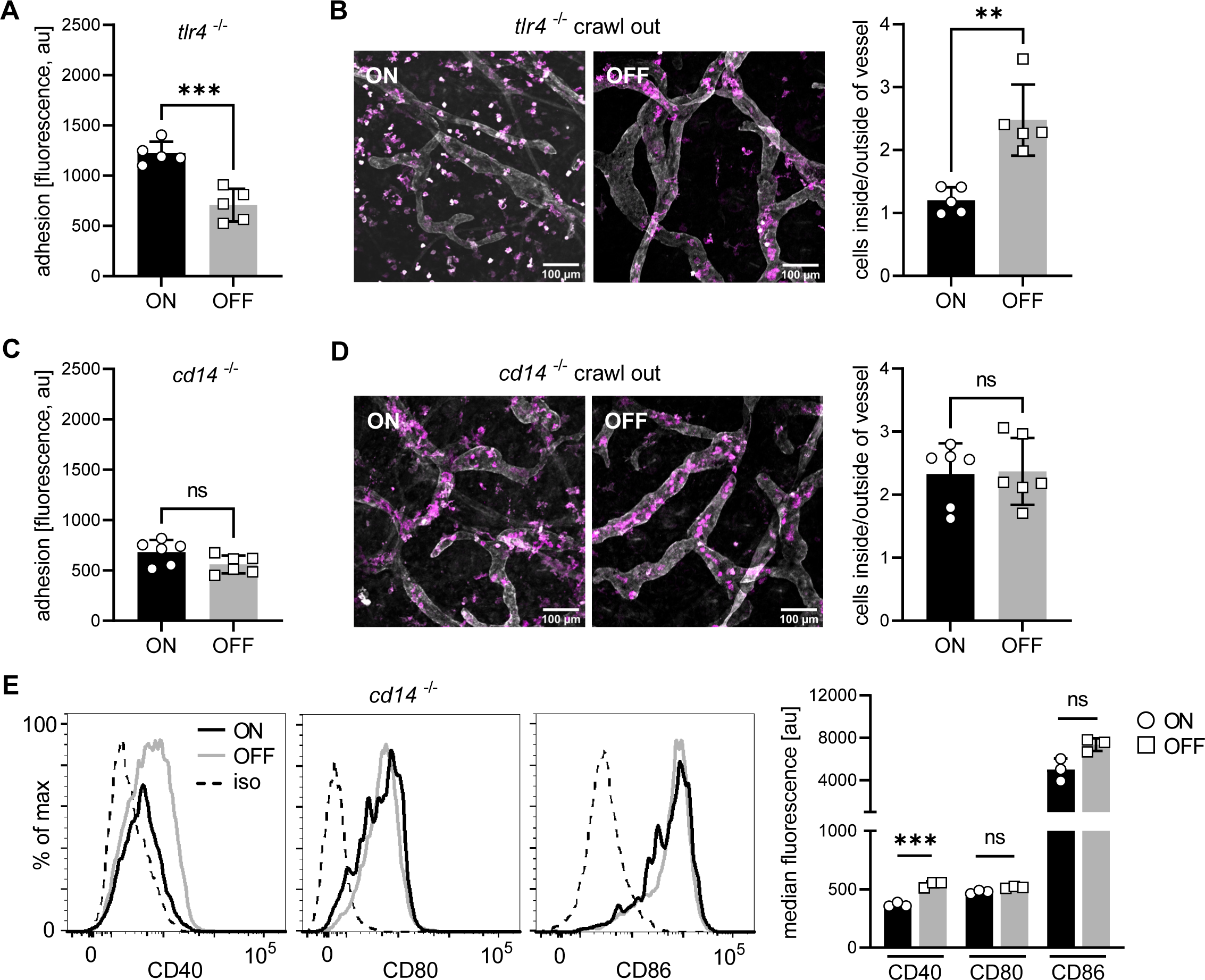
Interaction of type 1 piliated UPEC with CD14, but not TLR4, is important for the observed phenotypes. A. Adhesion assay of *tlr4^-/-^* and *cd14^-/-^* DCs. Quantification of fluorescence signal after ON (black) and OFF (grey) stimulation (5 biological replicates). (see also Supplementary Figure 3A) B. Ear crawl out assay of *tlr4^-/-^* DCs after ON and OFF stimulation. Endogenous DCs stained with anti-MHCII (magenta). Lymph vessels stained with anti-LYVE-1 (white) (left panel). Quantification of cells inside over outside of lymph vessel after ON (black) and OFF (grey) stimulation *tlr4^-/-^* DCs (right panel) (2 biological replicates). C. Quantification of the adhesion assay of *cd14^-/-^* DCs after ON (black) and OFF (grey) stimulation (5 biological replicates). (see also Supplementary Figure 3B) D. Ear crawl out assay of *cd14^-/-^* DCs after ON and OFF stimulation. Endogenous DCs stained with anti-MHCII (magenta). Lymph vessels stained with anti-LYVE-1 (white) (left panel). Quantification of cells inside over outside of lymph vessel after ON (black) and OFF (grey) stimulation *cd14^-/-^* DCs (right panel) (3 biological replicates). E. Expression level of co-stimulatory molecules (CD40, CD80, CD86) of *cd14^-/-^* DCs after stimulation with ON (black) and OFF mutants (grey) (left panel; iso – isotype control). Quantification of median fluorescence values of co-stimulatory molecules (right panel) (3 biological replicates). ns, not significant, ** p<0.05, *** p<0.01 by Student’s t test; data are represented as means ± SD

We then focused on CD14, a GPI anchored glycoprotein and co-receptor of TLR4, since a strong correlation has been reported between CD14 expression, integrin activity, and cell adhesion (Wright *et al,* 1991). We therefore performed adhesion and crawl out migration assays stimulating *cd14^-/-^* DCs with UPEC ON and OFF mutants and found that behavior of DCs was fully restored (Figure 3C and 3D and Supplementary Figure 3B). *cd14^-/-^* DCs also showed almost full rescue of the levels of co-stimulatory molecules after ON stimulation, compared to OFF stimulation (Figure 3E). We conclude that CD14 is required for the increases in integrin activity and decreases in co-stimulatory molecule expression after stimulation with type 1 piliated UPECs.

We next asked how type 1 pili interact with the CD14 receptor. *In silico* protein-protein docking analysis predicted strong binding of -56.98 kcal/mole between FimH (PDB: 6GTV; (Sauer *et al,* 2019)) and CD14 (PDB: 1WWL; (Kim *et al,* 2005)) (Table S1; File S1), which is stronger than the -49.81 kcal/mole between FimH and TLR4 (PDB: 3VQ2; (Ohto *et al,* 2012)) or the -35.55 kcal/mole for CD48 (PDB: 2PTV; (Velikovsky *et al,* 2007)), another FimH receptor (McArdel *et al,* 2016) (Table S1). Binding sites in FimH were located in its N-terminal domain and in CD14 within the central region of the crescent shaped monomer (Figure 4A), in an area not involved in LPS binding (Table S2) (Kim et al. 2005). Since CD14 of mouse (PDB: 1WWL) and human (PDB: 4GLP; (Kelley *et al,* 2013)) have highly similar secondary structures, but differ in their amino acid sequence (Kelley *et al,* 2013), we performed docking analysis with human CD14. We found predicted strong FimH binding (-55.95 kcal/mole) to be conserved in similar regions of both proteins but mediated by different amino acids (Table S1 and Table S3). This indicates that the difference in amino acid sequence between mouse and human CD14 seems negligible for the otherwise conserved strategy of how FimH of type 1 pili binds to CD14 receptor.

**Figure 4:**
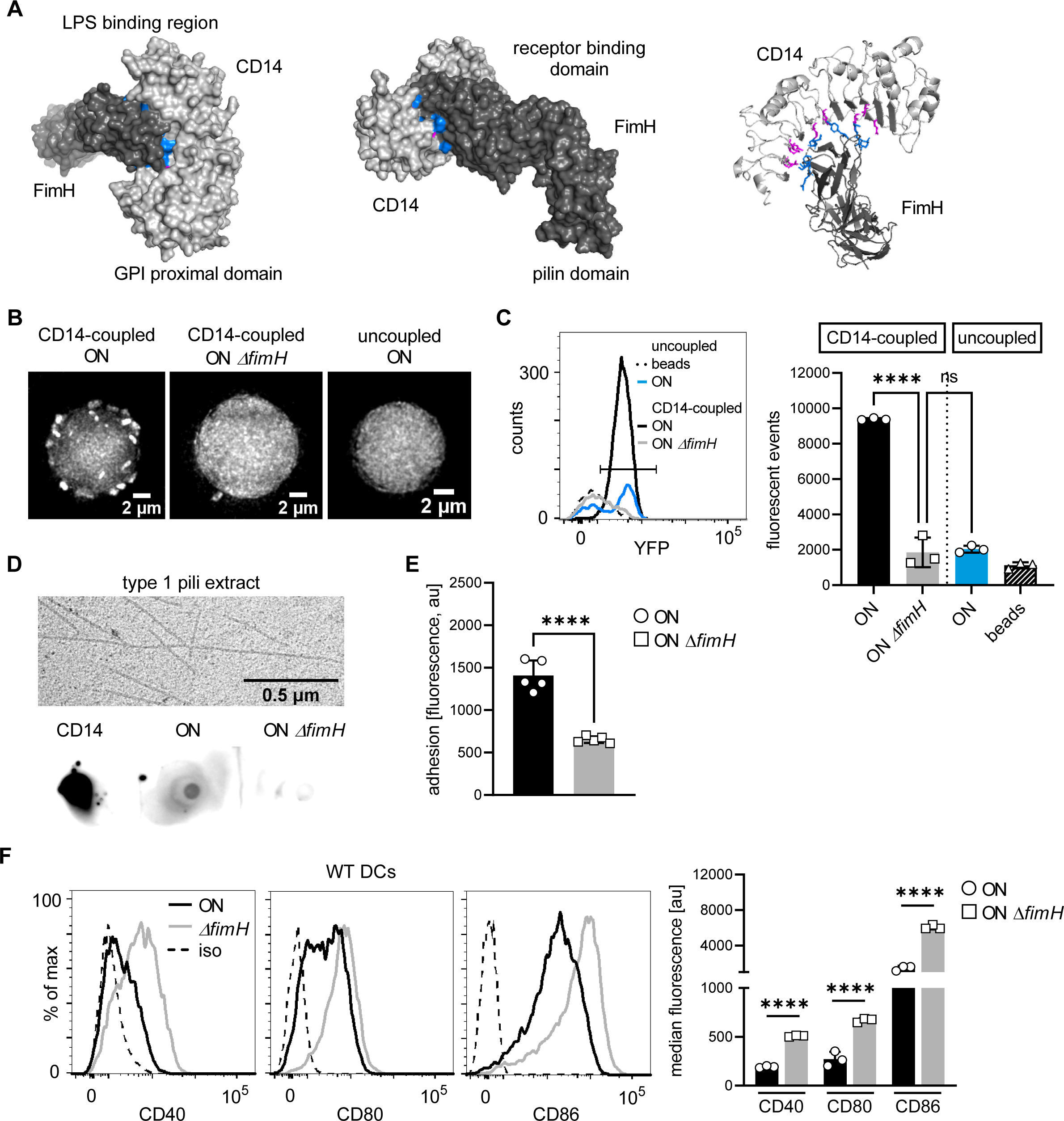
FimH binds to CD14 via protein-protein interactions and deletion of *fimH* **rescues adhesion and expression of co-stimulatory molecules.** A. In silico protein-protein docking analysis for FimH and CD14 (see also file S1 and Table S1). FimH is shown in dark grey and CD14 in light grey. Left panel shows surface plot of docked proteins from the view of CD14 on the membrane of the cell. Middle panel shows surface plot of docked proteins from the view of FimH on the membrane of *E. coli*. GPI proximal domain, LPS-binding, lectin and pilin domains are indicated. Right panel shows secondary structures of proteins. Top 10 amino acids predicted to interact during the binding are highlighted in blue for FimH and magenta for CD14. B. Microscopy images of bead binding assay. ON or ON *ΔfimH* mutants expressing a constitutive *yfp* fluorescent marker bound to CD14-coupled or uncoupled beads are shown. C. Flow cytometry analysis of bead binding assay. Histogram of fluorescence events in the bacteria gate (left panel). Uncoupled beads (dashed black), ON mutants bound to uncoupled beads (blue), ON mutants bound to CD14-coupled beads (black), ON *ΔfimH* bound to CD14-coupled beads (grey). Quantification of fluorescent events in the bacterial gate (right panel) (3 biological replicates). (see also Supplementary Figure 5) D. Type 1 pili extracts and dot blot assay. Electron microscopy images of type 1 pili extracts from ON and ON *ΔfimH* mutants (left panel). Dot blot assay of type 1 pili extracts with biotinylated CD14 (right panel). Pre-blotted biotinylated CD14 served as positive control. Bound CD14 was visualized with Streptavidin-HRP antibody. E. Adhesion assay of WT DCs after ON and ON *ΔfimH* stimulation. Quantification of fluorescence signals after ON (black) and ON *ΔfimH* (grey) stimulation (5 biological replicates). ON data are the same as in Figure 2C. (see also Supplementary Figure 3C) F. Expression level of co-stimulatory molecules (CD40, CD80, CD86) of WT DCs after stimulation with ON (black) and ON *ΔfimH* mutants (grey) (left panel; iso – isotype control). Quantification of median fluorescence values of co-stimulatory molecules (right panel) (3 biological replicates). ON data are the same as in Figure 1G. ns, not significant, **** p<0.001 by one-way ANOVA followed by Dunnett’s multiple comparisons (C) and by Student’s t test (E, F); data are represented as means ± SD

**Table 1.**
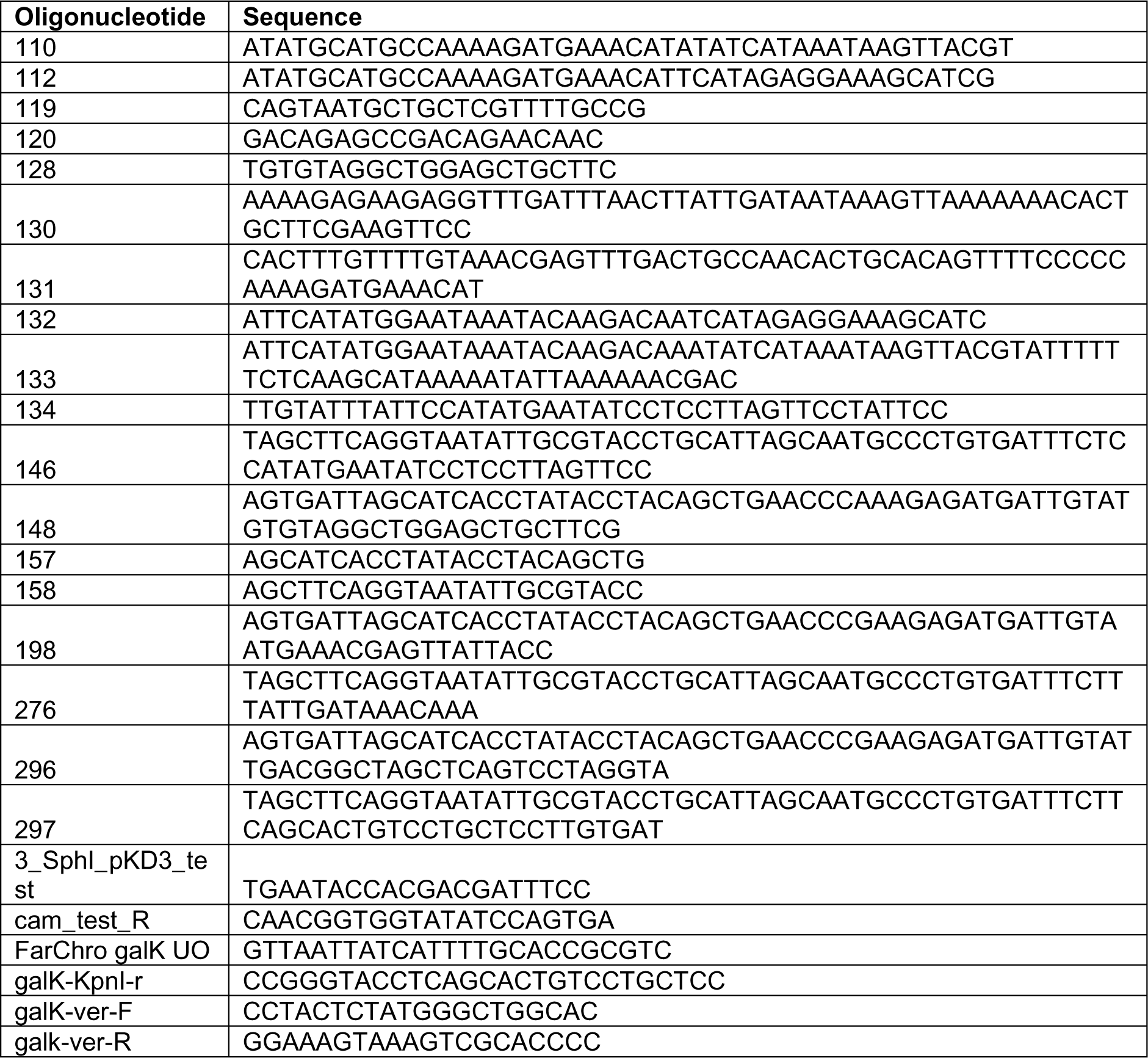
Primers used for cloning.

**Table 2.** Used strains.

| Strain | Reference |
| --- | --- |
| *E. coli* W (ATCC 9637) | Archer et al 2011 |
| CFT073 O6:K2:H1 (ATCC 700928) | Welch et al 2002 |
| KT179 (CFT073 locked-ON::FRT) | this paper |
| KT180 (CFT073 locked-OFF::FRT) | this paper |
| KT193 (CFT073 locked-ON::FRT Δ*fimH*::FRT) | this paper |
| KT232 (*E. coli* W locked-ON::FRT) | this paper |
| KT257 (CFT073 locked-ON::FRT Δ*fimH* attλ *P<sub>R</sub>-mVenus*::FRT) | this paper |
| KT260 (CFT073 locked-ON::FRT Δ*galK*::FRT *fimH*::*fimH*<sub>Y48A R98A T99A</sub>) | this paper |
| KT261 (CFT073 locked-ON::FRT Δ*galK*::FRT *fimH*::*fimH*<sub>Y48A</sub>) | this paper |
| KT262 (CFT073 locked-ON::FRT Δ*galK*::FRT *fimH*::*fimH*<sub>R98A</sub>) | this paper |
| KT263 (CFT073 locked-ON::FRT Δ*galK*::FRT *fimH*::*fimH*<sub>T99A</sub>) | this paper |
| MG002 (*E. coli* W locked-ON::FRT Δ*galK*::FRT *fimH*::*fimH*<sub>CFT073</sub>) | this paper |
| VG003 (CFT073 locked-ON::FRT attλ *P<sub>R</sub>-mVenus*::FRT) | this paper |

**Table 3.** Antibodies used.

| Antibody | Source | Identifier |
| --- | --- | --- |
| Rat anti-MHC II (I-A/I-E) eFluor450 (clone M5/114.15.2) | eBioscience | 48-5321-82 |
| Armenian hamster anti-CD11c APC (clone N418) | eBioscience | 17-0114-82 |
| Rat anti-CD18 PE (clone C71/16) | BD | 553293 |
| Rat anti-CD11b FITC (clone M1/70) | eBioscience | 11-0112-82 |
| Mouse anti-human CD11b (active epitope) APC (clone CBRM1/5) | eBioscience | 17-0113-41 |
| Rat anti-CD86 biotin (clone GL1) | eBioscience | 13-0862-85 |
| Armenian hamster anti-CD80 (clone 16-10A1) | eBioscience | 13-0801-85 |
| Rat anti-CD40 (clone 1C10) | Biolegend | 102802 |
| Armenian Hamster anti-CD103 Brilliant Violet 421 (clone 2E7) | Biolegend | 121421 |
| Rat anti-CD14 Brilliant Violet 421 (clone Sa14-2) | Biolegend | 123329 |
| Rat anti-CD4 eFluor450 (clone GK1.5) | eBioscience | 48-0042-82 |
| Armenian Hamster anti-CD69 APC-eFluor780 (clone H1.2F3) | eBioscience | 47-0691-82 |
| Rat anti-CD62-L PE (clone MEL-14) | eBioscience | 12-0621-82 |
| Rat IgG2b kappa Isotype Control eFluor450 (clone eB146/10H5) | eBioscience | 48-4031-82 |
| Armenian Hamster IgG Isotype Control APC (clone eBio299Arm) | eBioscience | 14-4888-81 |
| Rat IgG2a kappa Isotype Control PE (clone eBR2a) | eBioscience | 12-4321-81 |
| Rat IgG2b kappa Isotype Control FITC (clone eB149/10H5) | eBioscience | 11-4031-82 |
| Rat IgG1 kappa Isotype Control APC (clone eBRG1) | eBioscience | 17-4301-82 |
| Armenian Hamster IgG Brilliant Violet 421 (clone HTK888) | Biolegend | 400936 |
| Rat IgG2a kappa Isotype Control Biotin (clone RTK2758) | Stemcell Technologies | 60076 |
| Armenian Hamster IgG Isotype Control Biotin (clone eBio299Arm) | eBioscience | 13-4888-81 |
| Rat IgG2a kappa Isotype Control (clone RTK2758) | Stemcell Technologies | 60076 |
| Alexa Fluor 647-conjugated Streptavidin | Jackson ImmunoResearch | 016-600-084 |
| Donkey anti-rat IgG H+L Alexa Fluor488 AffiniPure F(ab')<sub>2</sub> Fragment | Jackson ImmunoResearch | 712-546-150 |
| Goat anti-rabbit Alexa Fluor488 | Invitrogen | A11008 |
| Rat anti-CD29 (clone 9EG7) | BD | 553715 |
| Armenian Hamster anti-CD29 PE (clone HMβ1-1) | Biolegend | 102207 |
| Armenian Hamster IgG Isotype Control PE (clone HTK888) | Biolegend | 400907 |
| M14-23 (anti-mouse CD14 antibody) | Biolegend | 150102 |
| Strep-Tactin HRP | iba-lifesciences | 2-1502-001 |
| Fc receptor block | eBioscience | 14-9161-73 |

To verify the predicted binding of FimH to CD14 *in vitro*, we generated UPEC ON mutants lacking the *fimH* gene (ON *ΔfimH*). Compared to *Salmonella*, where a *fimH* is necessary for the biosynthesis of type 1 fimbriae (Zeiner et al., 2012), *E. coli* mutants lacking *fimH* still express type 1 pili which are, however, non-functional (Maurer and Orndorff, 1985; Klemm and Christiansen, 1987; Jones et al., 1995). We confirmed that our ON *ΔfimH* mutants still express type 1 pili but lack the ability to agglutinate with yeast (Supplementary Figure 4A). We performed an immunoprecipitation-type approach using magnetic Protein A beads, a CD14-Fc chimeric protein and the bacterial mutants. First, the CD14-Fc chimera was coupled to the magnetic Protein A beads (Supplementary Figure 5A). Next, we introduced a constitutively expressed *yfp* fluorescent marker into the chromosome of the UPEC ON and ON *ΔfimH* mutants for tracking. Using fluorescence microscopy, we found abundant ON mutants bound per CD14-coupled bead, whereas binding of the *fimH* deletion mutants was scarce (Figure 4B). Binding of ON mutants was not due to unspecific binding of FimH to the bead matrix, as uncoupled beads also showed very scarce binding. Additionally, we analyzed the binding of bacteria to beads by flow cytometry by gating on the size parameters and fluorescence signal of the bacteria event population (see Supplementary Figure 5B for gating strategy and methods section for further technical details). ON mutants showed increased binding to CD14-coupled beads, when compared to the *fimH* deletion mutants or to ON mutants binding to uncoupled beads (Figure 4C). To further test if only FimH is necessary for binding to CD14, and not interactions with LPS on the bacterial membrane, we extracted type 1 pili (Sheikh *et al,* 2017) from UPEC ON and ON *ΔfimH* mutants and performed a dot blot assay using biotinylated CD14 and Streptavidin-HRP (Figure 4D). Biotinylated CD14 only bound to type 1 pili extracts from ON mutants, whereas no binding was observed to type 1 pili extracts from ON mutants lacking the FimH protein.

We next tested if binding of FimH to CD14 is causative for the increase in adhesion of DCs and the decrease in co-stimulator molecules on their membrane after UPEC ON stimulation. Indeed, deleting *fimH* from UPEC ON mutants fully rescued the adhesion and the expression of all co-stimulatory molecules when compared to stimulation with the UPEC ON mutants (Figure 4E and 4F).

Finally, we asked if expression of pathogenic FimH in an otherwise non-pathogenic bacterial background is sufficient to affect expression of co-stimulatory molecules on DCs. We constructed a locked-ON mutant of *E. coli* W, a non-pathogenic *E. coli* strain, and replaced the endogenous *fimH* gene with the pathogenic *fimH* variant of UPEC CFT073 strain. This non-pathogenic ON *fimH_CFT073_* mutant was able to agglutinate with yeast, confirming the correct insertion of the pathogenic *fimH* gene (Supplementary Figure 4B). However, expression of co-stimulatory molecules on DCs after stimulation with non-pathogenic *E. coli* W ON mutants or the non-pathogenic ON *fimH_CFT073_* mutant was barely affected (Supplementary Figure 4C).

Based on these experimental observations, we propose that CD14 is a novel target for the FimH protein of type 1 pili and that direct protein-protein interaction is the underlying mechanism of binding. FimH is necessary for the modulatory effects seen on DCs after stimulation with UPEC ON mutants, whereas expressing the pathogenic *fimH* gene alone is not sufficient to cause these effects in an otherwise non-pathogenic background.

### FimH amino acids predicted to bind are highly conserved and are partially located in the mannose-binding domain

The two-domain FimH protein consists of a receptor-binding domain and a pili-binding domain (Choudhury *et al,* 1999). The receptor-binding domain not only interacts with host receptors, like uroplakin Ia on bladder epithelial cells, but also highly specifically binds D-mannose which due to this specific interaction has been used in the treatment of UTIs (Wiles *et al,* 2008). The amino acids responsible for binding mannose are located in the mannose-binding pocket (P1, N46, D47, D54, Q133, N135, D140 and F142; Table S2) (Hung *et al,* 2002) and the tyrosine gate (Y48, I52, Y137; Table S2) (Touaibia *et al,* 2017). These residues are highly conserved among pathogenic and non-pathogenic *E. coli* strains (Figure 5A), whereas other amino acids that affect the flexibility of the FimH protein and therefore facilitate host colonization were found to be mutated in pathogenic *E. coli* (Chen *et al,* 2009; Sokurenko *et al,* 1998; Kalas *et al,* 2017). Interestingly, the most important amino acids of FimH we predicted to be responsible for binding to CD14 are highly conserved among several different *E. coli* strains, whether they are pathogenic or not (Figure 5A).

**Figure 5:**
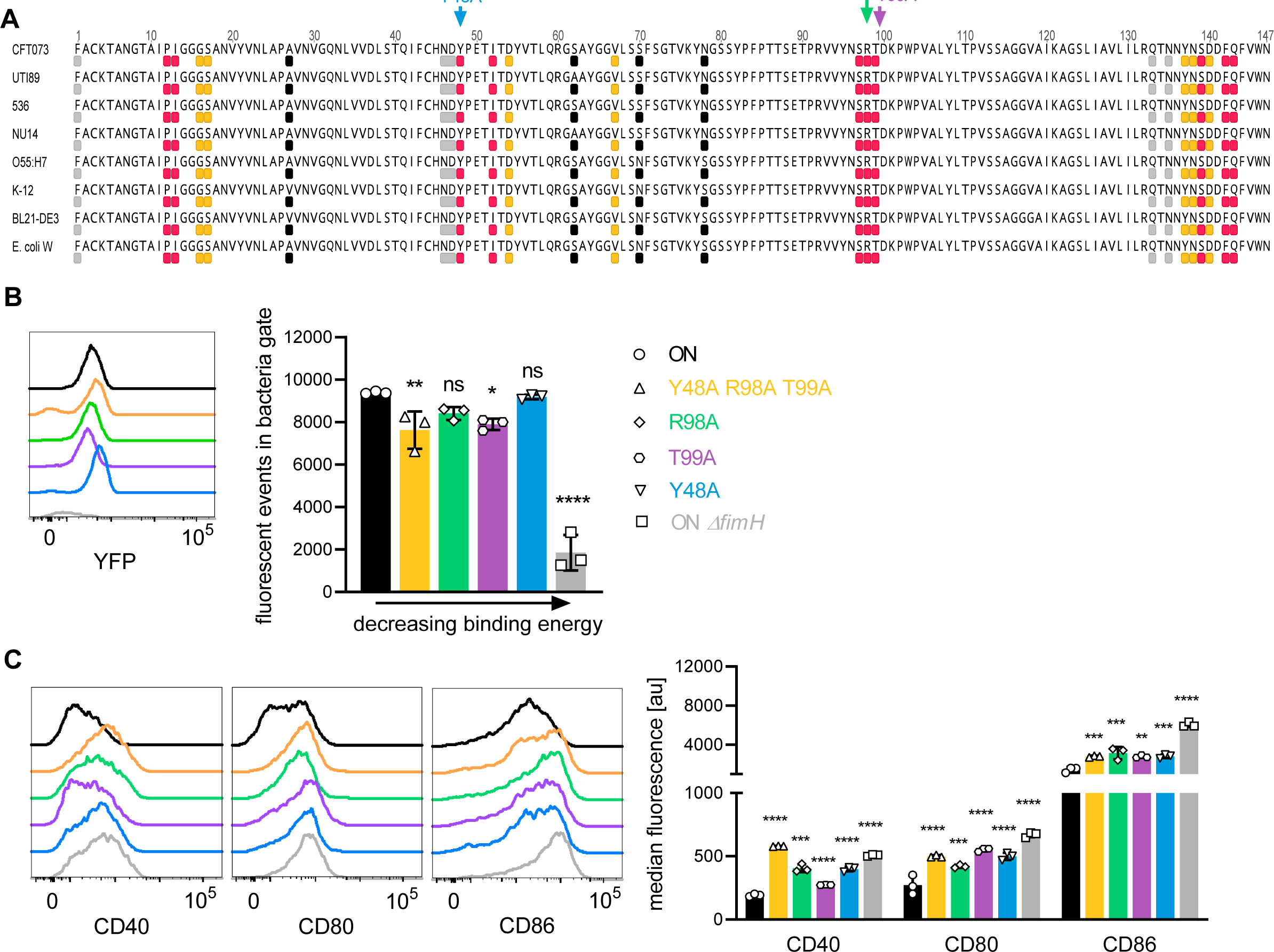
FimH binds to CD14 via highly conserved amino acid residues partially located in the mannose-binding domain. A. Receptor-binding domain of FimH from different *E. coli* strains is shown (UPEC: CFT073, UTI89, 536, NU14, EPEC: O55:H7; non-pathogenic; K-12, BL21-DE3, *E. coli* W). The top 10 amino acids on FimH showing strongest binding energy towards mouse CD14 (PDB: 1WWL) are shown in pink and towards human CD14 (PDB: 4GLP) are shown in orange. Amino acids I13, P12 and F42 are involved in both, mouse and human CD14, and therefore only shown in pink. Amino acids located in mannose-binding pocket and tyrosine gate are shown in grey. Amino acids I13, Y48, I52, Y137 and F142 are involved in mannose and CD14 binding and therefore only shown in pink. Common pathoadaptive mutations that differ between UPEC and non-pathogenic *E. coli* are shown in black. Amino acids mutated to generate FimH amino acid mutants were Y48 (binding energy -4.29 kcal/mole), T99 (binding energy -4.92 kcal/mole) and R98 (binding energy -7.23 kcal/mole) (see also Table S1 and S3). B. Bead binding assay of FimH amino acid mutants. Overlaid fluorescence events in the bacteria gate (left panel). ON (black), ON *ΔfimH* (grey), Y48A (blue), T99A (violet), R98A (green) and Y48A R98A T99A (yellow). Quantification of fluorescent events in the bacterial gate (right panel) (3 biological replicates). ON and ON *ΔfimH* data are the same as in Figure 4C. C. Overlaid expression levels of co-stimulatory molecules (CD40, CD80, CD86) of WT DCs after stimulation with ON (black), ON *ΔfimH* mutants (grey), and FimH amino acid mutants Y48A (blue), T99A (violet), R98A (green) and Y48A R98A T99A (yellow) (left panel). Quantification of median fluorescence values of co-stimulatory molecules (right panel) (3 biological replicates). ON data are the same as in Figure 1G. ns, not significant, * p<0.1, ** p<0.05, *** p<0.01, **** p<0.001 by one-way ANOVA followed by Dunnett’s multiple comparisons (the mean of the data were compared to the mean of ON); data are represented as means ± SD

To verify their significance for binding to CD14, we introduced mutations into the three most important amino acids. We exchanged amino acids R98 (binding energy -7.23 kcal/mole), T99 (binding energy -4.92 kcal/mole) and Y48 (binding energy -4.29 kcal/mole) individually to alanine or all three at the same time creating a triple mutant. All four FimH mutants were still able to bind to CD14-coupled beads and only the triple mutant showed some decrease in binding (Figure 5B). Stimulating DCs with the FimH amino acid mutants showed partial rescue for co-stimulatory molecule expression levels (Figure 5C). This suggests that not only these individual amino acids mediate binding to CD14, but most likely several other amino acids and thus the supporting secondary structure of the FimH protein are involved.

Since Y48 and other identified FimH amino acids with weaker binding energies for CD14 (Table S2) are located in the mannose binding pocket, we were interested whether FimH antagonists, such as D-mannose or the low molecular weight mannose derivate M4284 (Schönemann *et al,* 2019), disrupt the interaction. Additionally, we tested a blocking CD14 antibody (M14-23) (Tsukamoto *et al,* 2010) for its ability to inhibit binding of FimH to CD14. Our experiments showed that 175 µM D-mannose was not sufficient to block binding of UPEC ON bacteria to CD14, whereas 1 mM D-mannose and M4284 at 10 µM were able to inhibit binding of ON mutants to CD14-coupled beads (Figure 6A). The blocking CD14 antibody reduced bacteria binding to beads by roughly 25 % but was the most effective at rescuing expression of co-stimulatory molecules on DCs among all tested components (Figure 6B). Thus, existing FimH antagonists as well as blocking CD14 antibodies show certain potential to treat recurring or persistent infections caused by type 1 piliated pathogens.

**Figure 6:**
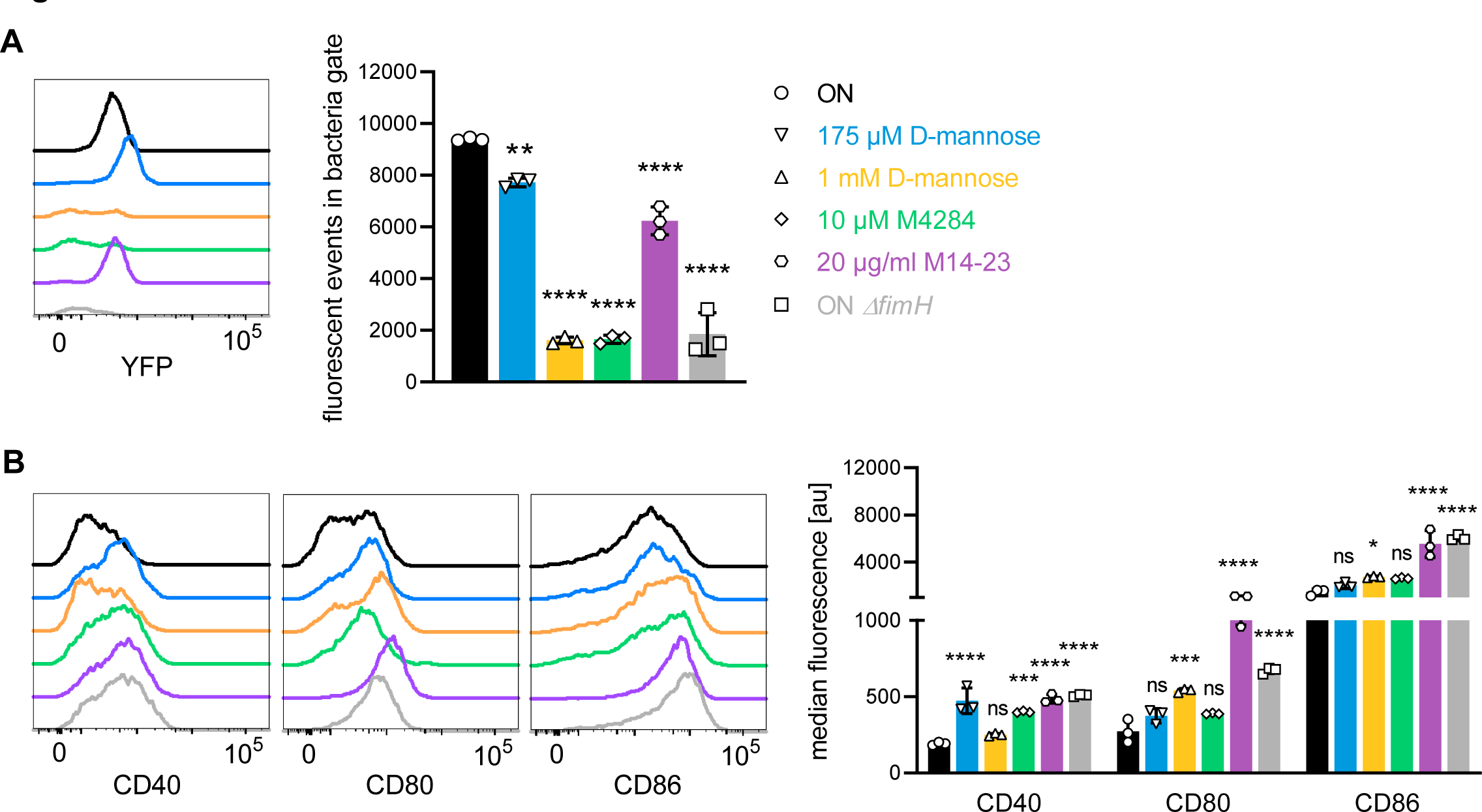
FimH antagonists and a blocking CD14 antibody (partially) block binding and rescue expression of co-stimulatory molecules on DCs. A. Bead binding assay of ON mutants in the presence of FimH antagonists and blocking CD14 antibody. Overlaid fluorescence events in the bacteria gate (left panel). ON (black), ON *ΔfimH* mutants (grey), 175 µM D-mannose (blue), 1 mM D-mannose (yellow), 10 µM M4284 (green) and 20 µg/ml M14-23 antibody (violet). Quantification of fluorescent events in the bacterial gate (right panel) (3 biological replicates). B. Overlaid expression levels of co-stimulatory molecules (CD40, CD80, CD86) of WT DCs after stimulation with ON (black), ON *ΔfimH* mutants (grey), and ON stimulation in presence of 175 µM D-mannose (blue), 1 mM D-mannose (yellow), 10 µM M4284 (green) and 20 µg/ml M14-23 antibody (violet) (left panel). Quantification of median fluorescence values of co-stimulatory molecules (right panel) (3 biological replicates). ns, not significant, * p<0.1, ** p<0.05, *** p<0.01, **** p<0.001 by one-way ANOVA followed by Dunnett’s multiple comparisons (the mean of the data were compared to the mean of ON); data are represented as means ± SD

## Discussion

Here, we uncovered that type 1 piliated UPECs target the CD14 glycoprotein on the surface of DCs to shut down the migratory capacity of DCs and their ability to interact with and activate T cells.

It has been proposed that the tight regulation of UPECs type 1 pili expression has evolved to limit exposure to the host immune system, allowing the pathogen to establish a persistent infection (Donnenberg, 2013). In this study we uncovered and characterized a different fundamental role of type 1 pili, as modulators of the innate and adaptive immune response. We found that type 1 piliated UPECs decreased the migratory capacity of DCs by increasing their adhesion to other cells, such as T cells, and the extracellular matrix by over-activation of integrins. The effective and timely transition from adhesion to the migration phenotype is essential for DCs to migrate from the site of the actual infection to the lymph node in order to interact with lymphocytes. To achieve these migratory and signaling tasks, DCs need to dynamically regulate integrin mediated adhesion, and we found that over-activation of integrins triggered by type 1 piliated UPECs leads to effective immobilization to the extracellular matrix and decreased turnover of cell-cell interactions. Over-activation of integrins, by hijacking integrin-linked kinases leading to decreased turnover of focal adhesions, was already shown to subvert innate immunity during *Shigella* and enterohemorrhagic *E. coli* (EHEC) infections (Kim *et al,* 2009; Shames *et al,* 2010), however UPECs do not express the responsible genes. Moreover, we found that type 1 piliated UPECs also decreased the expression of co-stimulatory molecules, which are essential for T cell activation and proliferation.

We identified CD14, a GPI-anchored glycoprotein, as the direct target for the FimH protein of type 1 pili. In analogy, a previous study showed that the *fimA*-encoded major type V fimbriae of the oral pathogen *Porphyromonas gingivalis* also bind CD14 and thereby increase integrin-mediated adhesion by activating CD11b integrin (Harokopakis & Hajishengallis, 2005). Using in silico protein-protein docking analysis we found strong predicted binding between CD14 and FimH through specific amino acids in both proteins. Interestingly, the respective FimH amino acids are highly conserved not only among pathogenic, but also non-pathogenic *E. coli* strains. However, although we found FimH to be necessary, we did not find the simple presence of pathogenic FimH in an otherwise non-pathogenic genetic background to be sufficient for suppression of the immune response. We therefore hypothesize that the predicted amino acids are mainly important for the pathogens to interact with and thereby to dampen the activation of immune cells and thus the immune response in the host to persist “commensally” for prolonged times. However, it is unknown which other gene(s) work together with *fimH*, as the UPEC strain CFT073 carries accessory genes encoded by 13 pathogenicity islands (Lloyd *et al,* 2007). Generally, the pathogenic potential of UPECs does not seem to be the result of a defined virulence gene, but rather a combination of effects by several genes (Touchon *et al,* 2009).

Given how pathogenic bacteria circumvent the host immune response by generating protected niches (Grant & Hung, 2013) and the constantly growing presence of multi-drug resistant strains (Boucher *et al,* 2009), treatment of persistent or recurring infections is increasingly challenging. Although vaccination approaches against FimH (Eldridge *et al,* 2020) or non-conventional treatments, such as small mannosides (Schönemann *et al,* 2019; Mydock-McGrane *et al,* 2016), showed promising results, treatments of infections caused by type 1 piliated pathogens remains difficult. Our findings underscore the importance of the mannose-binding domain, and several conserved amino acid residues in this domain, in the interaction between FimH and CD14. For example, R98, showing the strongest binding to CD14 in our predictions, was identified to be important to stabilize the protein-ligand interaction (Han *et al,* 2010). However, targeting this residue only, was not sufficient to boost affinity of FimH antagonists (Tomašič *et al,* 2021). Given our results, we hypothesize that this could be because not individual amino acids, but the overall secondary structure of the mannose-binding domain of FimH is important for the interaction with host receptors. Blocking CD14 antibodies, such as the human anti-CD14 antibody IC14 (Axtelle & Pribble, 2001), could represent an alternative way to treat recurring infections caused by type 1 piliiated pathogens, such as UTIs or inflammatory bowel diseases like Crohn’s disease (Sivignon *et al,* 2017), given that CD14 is expressed by hematopoietic but also non-hematopoietic cells (Zanoni & Granucci, 2013).

Although opportunistic pathogens continuously circulate in humans as asymptomatic colonizers, they occasionally cause symptomatic acute or even chronic infections in some individuals (Donnenberg, 2013). Opportunistic pathogens use type 1 pili to adhere to and invade into host epithelial cells. While the cellular invasion represents a *de facto* passive mechanism for the pathogens to hide from the immune system and to establish persistent or recurring infections, we found that UPECs use type 1 pili to actively manipulate the behavior of innate immune cells, and thus also the adaptive immune response, by direct interaction of FimH and CD14 receptor on DCs. Since CD14 is a multi-functional co-receptor of several immune cell types (Zanoni & Granucci, 2013) and type 1 pili are expressed not only by pathogenic but also by non-pathogenic *E. coli* (Shawki & McCole, 2017; Croxen & Finlay, 2010), our findings add a new layer of complexity to the physiological relevance of type 1 pili – as modulators of the immune response in general and specifically during persistent and recurring infections.

## Supplementary Figure Legends

**Supplementary Figure 1:**
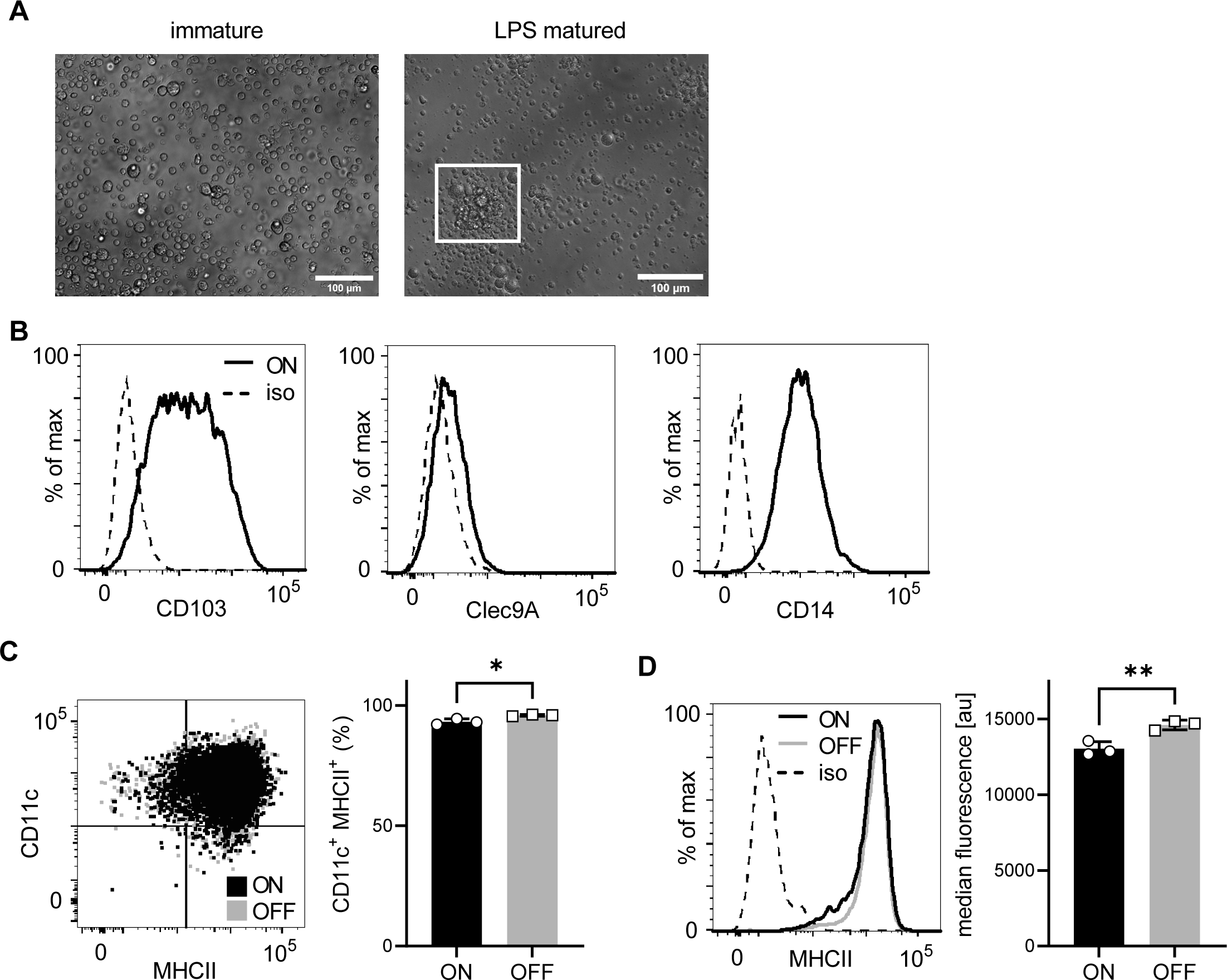
In vitro generated DCs from Hoxb8-progenitor cells resemble iCD103 DCs and differentiation of cells is not different after UPEC ON and OFF stimulation. A. Upon stimulation with recombinant LPS (200 ng/ml) immature DCs change to a mature phenotype and cluster together (see cells highlighted in inlay). B. In vitro generated iCD103 DCs express CD103, Clec9A and CD14. C. Differentiated DCs after ON (black) and OFF (grey) stimulation are defined by expression of CD11c integrin (DC marker) and upregulation of MHCII (activation marker) (left panel). Quantification of differentiated DCs after ON (black) and OFF (grey) stimulation (right panel) (3 biological replicates). D. Histograms of MHCII of WT DCs after stimulation with ON (black) and ON *ΔfimH* mutants (grey) (left panel; iso – isotype control). Quantification of median fluorescence values of MHCII (right panel) (3 biological replicates). ns, not significant, * p<0.1, ** p<0.05 by Student’s t test; data are represented as means ± SD

**Supplementary Figure 2:**
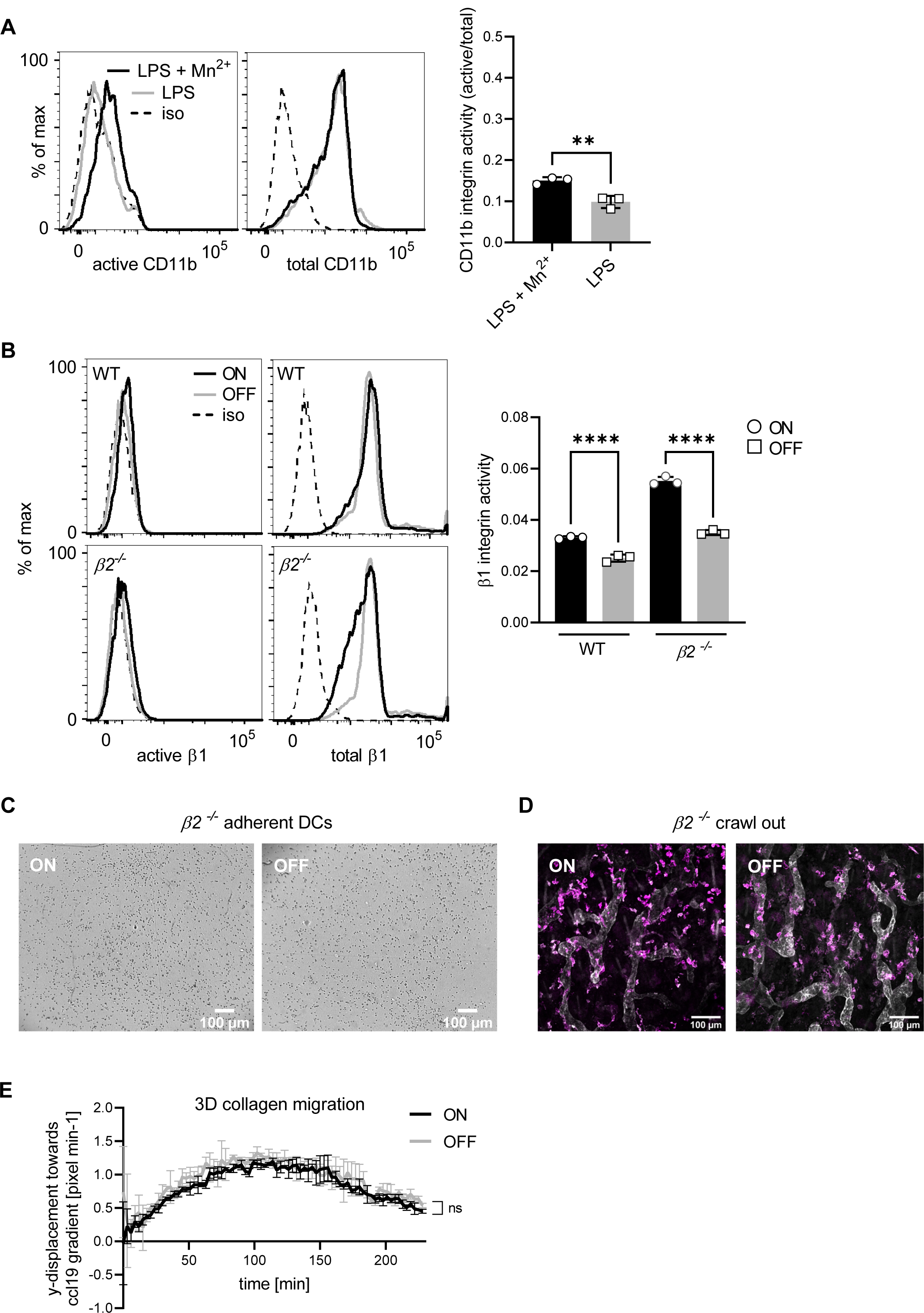
Type 1 piliated UPECs do not affect integrin-independent migration. A. Expression level of active and total CD11b integrin of WT DCs after LPS plus Mn^2+^ stimulation (black) and LPS only stimulation (grey) (left panel; iso – isotype control). Quantification of CD11b integrin activity (active/total levels of CD11b) (right panel) (3 biological replicates) (see also Figure 2B). B. Expression level of active and total β1 integrin after ON (black) and OFF (grey) stimulation of WT and *β2^-/-^* DCs (left panel; iso – isotype control). Quantification of β1 integrin activity (active/total levels of CD11b) (right panel) (3 biological replicates). It should be noted, that β1 integrin activity after UPEC ON stimulation of *β2^-/-^* DCs might be artificially increased due to a slight decrease in total β1 integrin staining after ON stimulation. C. Adhesion assay of *β2^-/-^* DCs after ON and OFF stimulation (quantification see Figure 2C). D. Ear crawl out assay of *β2^-/-^* DCs after ON and OFF stimulation. Endogenous DCs stained with anti-MHCII (magenta). Lymph vessels stained with anti-LYVE-1 (white) (left panel). Quantification of cells inside over outside of lymph vessel after ON (black) and OFF (grey) stimulation *β2^-/-^* DCs (right panel) (3 biological replicates) (quantification see Figure 2E). E. 3D collagen migration assay of WT DCs after stimulation with ON (black) and OFF (grey) mutants (3 biological replicates). ns, not significant, ** p<0.05, **** p<0.001 by one-way ANOVA followed by Dunnett’s multiple comparisons (D) and by Student’s t test (A and E); data are represented as means ± SD

**Supplementary Figure 3:**
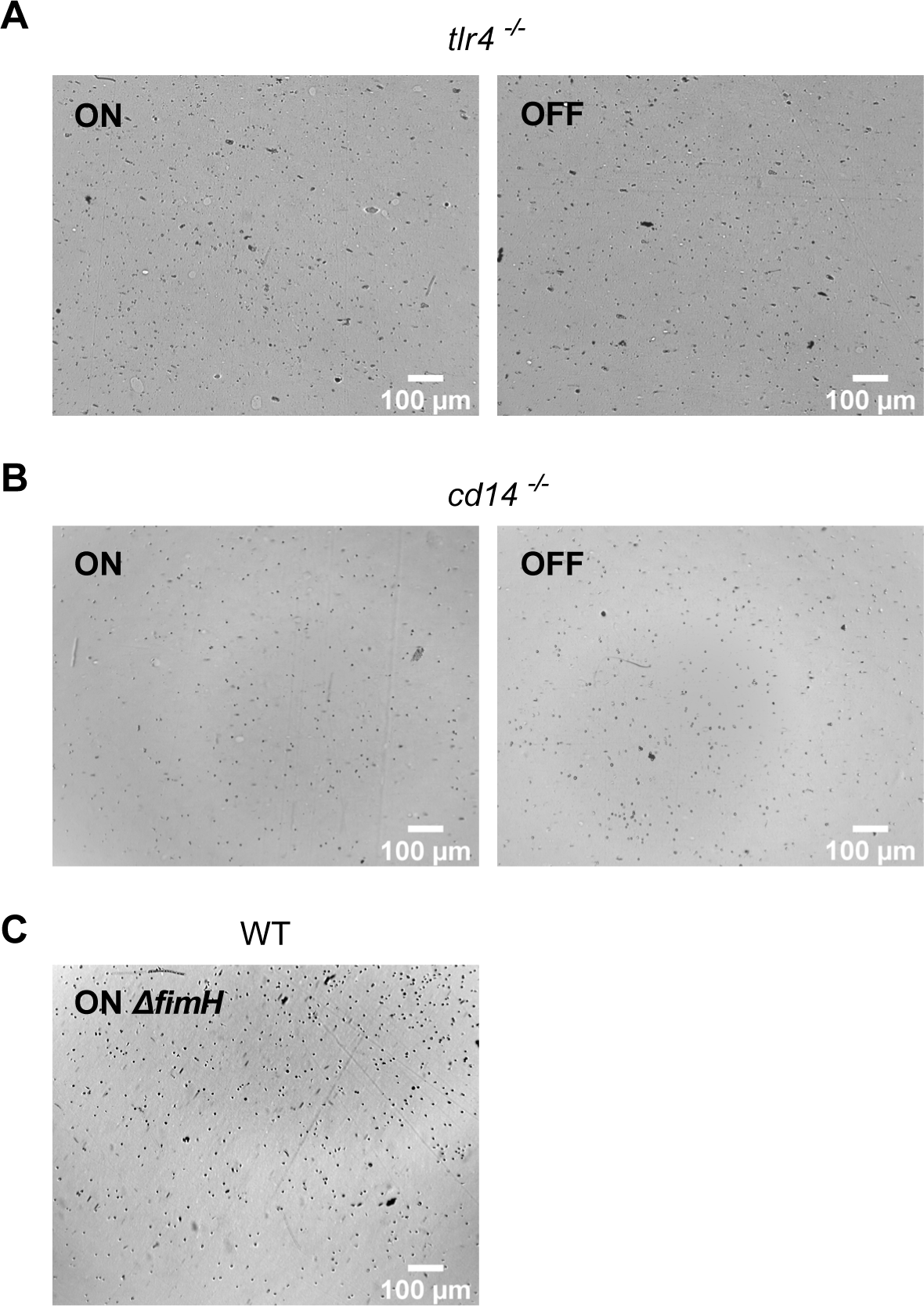
FimH and CD14, but not TLR4, are important for the observed adhesion phenotype. A. Adhesion assay of *tlr4^-/-^* DCs after ON and OFF stimulation (quantification see Figure 3A). B. Adhesion assay of *cd14^-/-^* DCs after ON and OFF stimulation (quantification see Figure 3B). C. Adhesion assay of WT DCs after ON *ΔfimH* stimulation (quantification see Figure 4E).

**Supplementary Figure 4:**
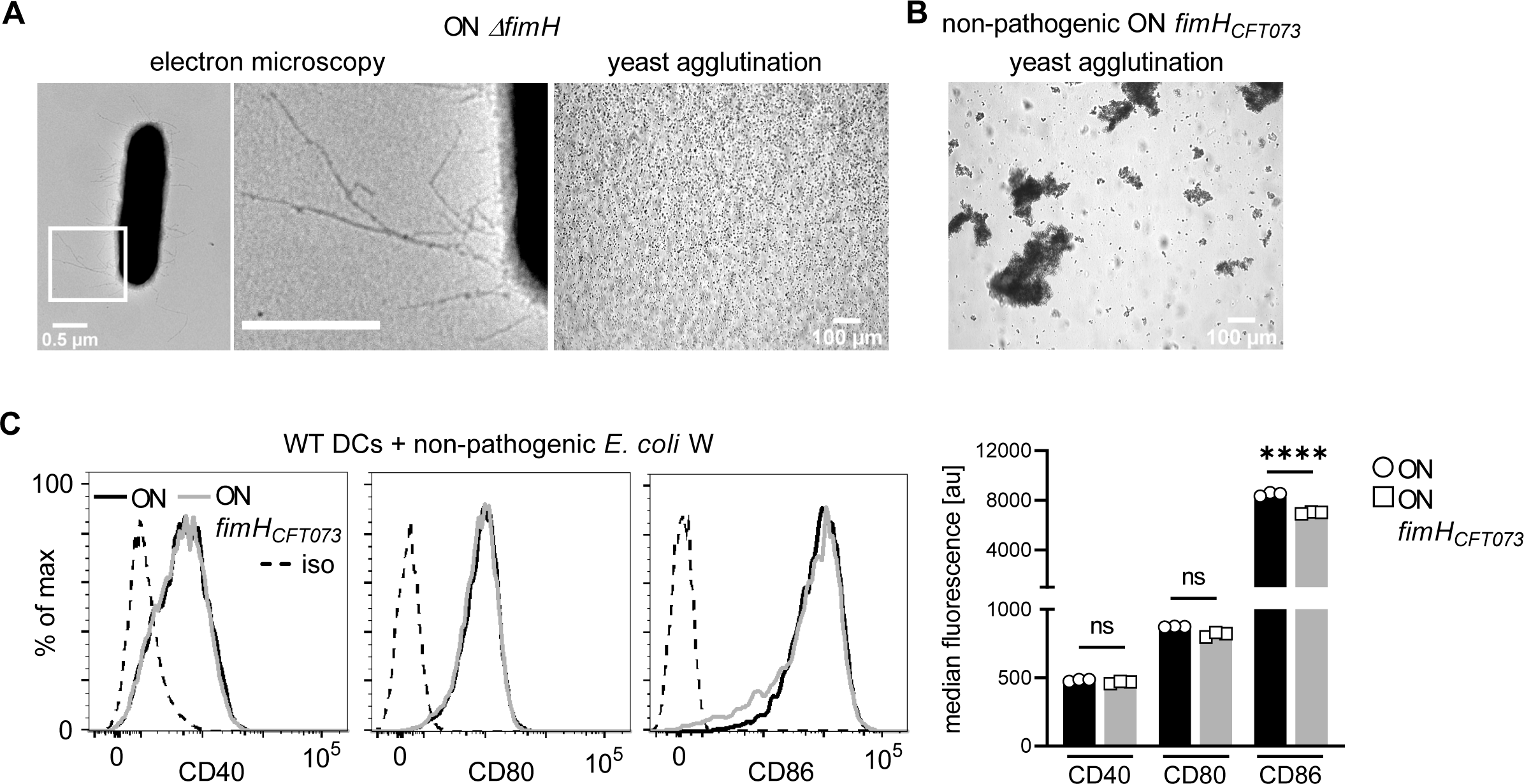
FimH is necessary, but it is not sufficient – bacteria need to have a pathogenic genetic background to cause adverse effects to DCs. A. Electron microscopy images, with zoomed in regions marked in inlays, (left panel) and yeast agglutination assay (right panel) of ON *ΔfimH* mutant. B. Yeast agglutination assay non-pathogenic ON mutant with pathogenic *fimH* (ON *fimH_CFT073_*). C. Expression levels of co-stimulatory molecules (CD40, CD80, CD86) of WT DCs after stimulation with non-pathogenic ON (black) and non-pathogenic ON *fimH_CFT073_* mutants (grey) (left panel; iso – isotype control). Quantification of median fluorescence values of co-stimulatory molecules (right panel) (3 biological replicates). ns, not significant, **** p<0.001 by Student’s t test; data are represented as means ± SD

**Supplementary Figure 5:**
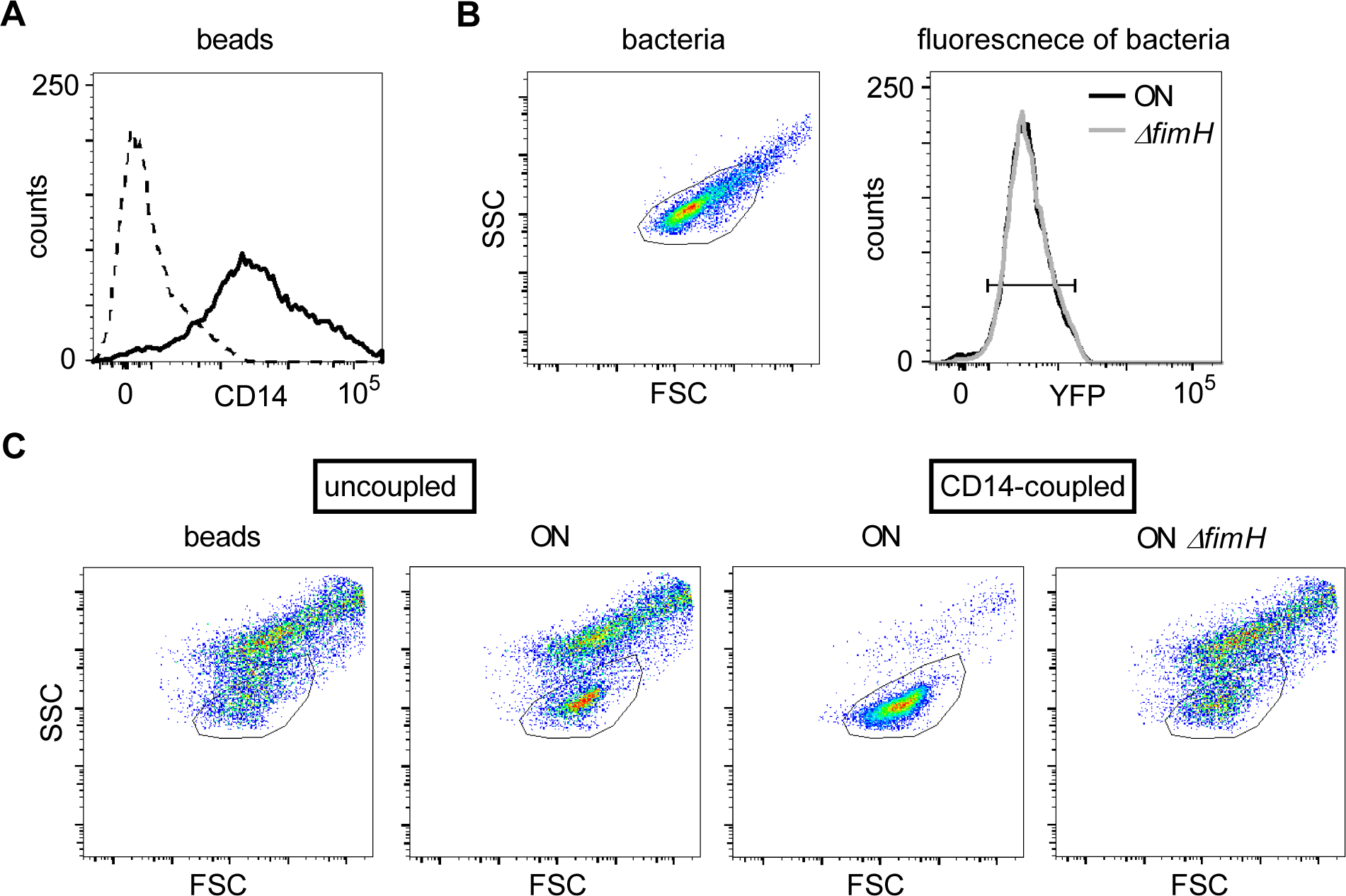
Bead assay using CD14-coupled magnetic beads. A. Coupling of recombinant CD14-Fc to the magnetic Protein A beads was confirmed by staining with anti-CD14 antibody (dashed black – uncoupled beads, solid black – CD14-coupled beads). B. The scatter properties for the bead assay were defined by FSC and SSC properties of the bacteria (left panel). The *yfp* fluorescence signal of ON (black) and ON *ΔfimH* (grey) mutants in the bacteria gate was used to define the fluorescent properties for analysis (right panel). C. Events (beads ± bacteria) in FSC and SSC were recorded and are shown as dot plots. The bacteria gate as defined in A is shown.

**Supplementary Figure 6:**
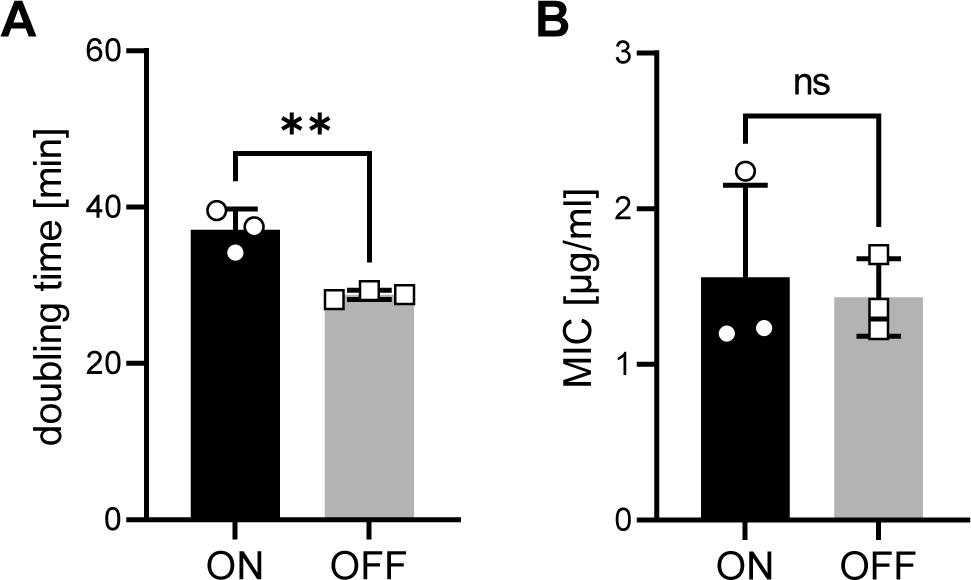
UPEC ON and OFF mutants exhibit slightly different growth rates but no difference in minimal inhibitory concentration (MIC) to gentamicin. A. Doubling time of ON and OFF mutants in R10H20. B. MIC to gentamicin for ON and OFF mutants (MIC = 1.5 µg/ml; 5x MIC was used in the infection assays after 1 h of co-culture of bacteria and DCs). ns, not significant, ** p<0.05 by Student’s t test; data are represented as means ± SD

## Methods

### Experimental Model and Subject Details

#### Animals

Mice were bred and maintained at the local animal facility in accordance with the IST Austria ethics commission. All experiments were conducted in accordance with the Austrian law for animal experiments. Permission (BMWFW-66.018/0010-WF/V/3b/2016) was granted by the Austrian federal ministry of science, research and economy.

#### Cell culture

R10 medium, RPMI 1640 + 10 % FCS, 2 mM L-Glutamine, 50 µM beta-mercaptoethanol, 100 U/ml Penicillin and 100 µg/ml Streptomycin, was used as basic medium. Stem cell medium was supplemented with 10 ng/ml IL-3, 20 ng/ml IL-6 and 1 % stem cell factor (SCF) supernatant produced by B16 melanoma cells. Hoxb8 medium was supplemented with 125 ng/ml Flt3 and 1 µM estradiol. iCD medium was supplemented with 2 ng/ml GM-CSF and 75 ng/ml Flt3. GM-CSF and Flt3 supernatants were produced by hybridoma cells and concentration of cytokines was measured by ELISA. All media were used pre-warmed. Cells were grown routinely at 37 °C with 5 % CO_2_. For infection and subsequent assays, cells were cultured in R10 medium without antibiotics and buffered with 20 mM Hepes (R10H20) in the absence of CO_2_.

#### WT, β2^-/-^, tlr4^-/-^ and cd14^-/-^ Hoxb8 cell generation

5-week-old wild type C57BL/6J, *β2^-/-^* (B6.129S7-*Itgb2^tm1Bay^*/J), *tlr4^-/-^* (B6(Cg)-*Tlr4*^tm1.2Karp^/J) and *cd14^-/-^* (B6.129S4-*Cd14*^tm1Frm^/J) mice were obtained from the Jackson Laboratory. Immortalization of bone marrow cells was performed as described previously (Leithner *et al,* 2018; Redecke *et al,* 2013). In brief, bone marrow was isolated from the femur and tibia by centrifugation and cells were precultured in stem cell medium for 3 days to enter the cell cycle. 1*10^5^ cells were spin-infected with Hoxb8-MSCV retrovirus using lipofectamine at 1000 g for 1 h in Hoxb8 medium. Cells were fed and split every few days for 3-4 weeks until all uninfected cells died off and only Hoxb8 infected, immortalized, cells were left.

#### Dendritic cell differentiation

iCD103 DCs, expressing CD103, were differentiated from Hoxb8 progenitor cells as described previously for bone marrow cells with minor modifications (Mayer *et al,* 2014). In brief, Hoxb8 cells were seeded at a density of 3*10^5^ cells into 10 cm bacterial culture dishes in 10 ml iCD medium. On day 3 cells were split 1:2 and toped up with fresh iCD medium to 10 ml. On day 6, cells were fed with 10 ml iCD medium and non-adherent iCD103 DCs were frozen on day 9. For images of immature and matured cells, as well as flow cytometry staining of different surface markers see Supplementary Figure 1.

Frozen DCs were allowed to recover after thawing for at least 4 h before infection. Non-adherent cells were counted and seeded at a density of 1-2*10^5^/ml in R10H20 medium. Assays were performed with DCs that were either stimulated with bacteria or recombinant LPS at 200 ng/ml. *β2^-/-^* DCs were purified from potential other cells using the EasySep^TM^ mouse Pan-DC enrichment kit (Stemcell Technologies) and allowed to rest for 1 h before the infection assay.

#### Bacterial strain construction

For cloning, strains were grown routinely in LB medium. Plasmids were maintained at 100 µg/ml ampicillin or 50 µg/ml kanamycin. Single copy integration was performed at 12 µg/ml chloramphenicol or 25 µg/ml kanamycin. For experiments, strains were grown at 37 °C in R10H20 medium without antibiotics or in medium containing 0.5 % casamino acids, 1x M9 salts, 1 mM MgSO4, 0.1 mM CaCl2 and 0.5 % glycerol (CAA M9 glycerol). Primers used for cloning are listed in Table 1 and strains used are listed in Table 2.

##### Locked mutants

*E. coli* W ((Archer *et al,* 2011); ATCC 9637) and clinical isolate CFT073 ((Welch *et al,* 2002); ATCC 7000928; a kind gift of Ulrich Dobrindt) were used as well as derivatives of those strains.

Phase-locked mutants were generated by replacing the 9 bp long recognition site for the site-specific recombinases FimB and FimE in the internal repeat region on the left site of the fim switch (*fimS)* with an FRT site (Figure 1C). The *fimS* region was amplified either in the ON (primers 132 and 110) or OFF (primers 133 and 112) orientation from the chromosome of the wild type strain but omitting the 9 bp recognition site on the left site. An FRT-removable chloramphenicol resistance marker was amplified from pKD3 plasmid using primers 128 and 134 (Datsenko & Wanner, 2000). The resistance marker was assembled left of the amplified *fimS* regions using NEBuilder assembly kit (NEB). The assembled DNA fragments were PCR amplified using primers 130 and 131 and integrated into the chromosome of the respective strains instead of the endogenous *fimS* using lambda red recombination (Datsenko & Wanner, 2000). In brief, *E. coli* W or CFT073 wild type bacteria were transformed with pSIM6 plasmid expressing thermal inducible Red genes under control of the native λ phage *pL* (Datta *et al,* 2006) and selected with ampicillin at 30 °C. After inducing expression of lambda red genes at 42 °C for 15 min, bacteria were made electrocompetent and transformed with 100 ng of the cleaned PCR fragments. Bacteria were allowed to recover in LB medium for 1 h at 37 °C before spreading on LB plates containing chloramphenicol.

After verifying single copy integration using primers 119, 120, cam_test_R and 3_SphI_pKD3_test, the resistance marker was subsequently removed using pCP20 plasmid (Cherepanov & Wackernagel, 1995). The mutated *fimS* region was sequenced to confirm deletion of the 9 bp long recombinase recognition site on the left site, but a fully intact recognition site on the right site. Presence or absence of the type 1 pilus on the bacterial outer membrane was confirmed by electron microscopy and yeast agglutination assay. Resulting locked mutants were: CFT073 locked-ON – KT179, CFT073 locked-OFF – KT180 and *E. coli* W locked-ON – KT232.

##### FimH deletion mutant

*fimH* gene from CFT073 locked-ON mutants (KT179) was deleted by lambda red recombination. The FRT-flanked chloramphenicol resistance marker from pKD3 plasmid was amplified using primers 146 and 148 and integrated into the chromosome of KT179 strain. Successful deletion was confirmed by PCR (primers 157 and 158). Resistance was flipped using pCP20 plasmid resulting in CFT073 locked-ON *ΔfimH* mutant – KT193. Presence of type 1 pili was confirmed by electron microscopy. Absence of *fimH* was confirmed by sequencing and yeast agglutination assay.

##### Chromosomal yfp marker

*mVenus* driven by the right site of the lambda PO was integrated in the lambda phage attachment site on the chromosome of CFT073 locked-ON (KT179) and CFT073 locked-ON *ΔfimH* (KT193) mutants using CRIM integration (Haldimann & Wanner, 2001). In brief, KT179 and KT193 strains were transformed with pInt-ts helper plasmid and selected on ampicillin plates at 30 °C. Bacteria were made electrocompetent and transformed with *PR-mVenus* carrying pAH120-frt-cat integration plasmid. After recovery in LB medium for 1 h at 37 °C, expression of lambda red genes was induced at 42 °C for 15 min before spreading on LB plates containing chloramphenicol.

Single copy integration of the CRIM plasmid was verified with PCR as mentioned previously (Haldimann & Wanner, 2001). Since pAH120-frt-cat was designed to have a FRT flanked chloramphenicol resistance marker (Nikolic *et al,* 2018), resistance was subsequently removed using pCP20 plasmid (Cherepanov & Wackernagel, 1995). Resulting mutants were: CFT073 locked-ON *attλ PR*-mVenus – VG003 and CFT073 locked-ON *ΔfimH attλ PR*-mVenus – KT257.

##### FimH replacement mutant

The endogenous *fimH* gene of the non-pathogenic *E. coli* W strain was exchanged scar-less with *fimH* of the pathogenic UPEC strain CFT073 using *galK* selection/counter-selection (Kavčič *et al,* 2020). The FRT-flanked kanamycin resistance marker from gDNA harboring *ΔgalK::kan* (gift of Bor Kavčič) was amplified using primers galK-ver-F and galK-ver-R (gift of Bor Kavčič) and integrated into the *galK* gene of *E. coli* W locked-ON (KT232). Loss of *galK* gene was confirmed by PCR using primers FarChro galK UO and galK-KpnI-r (gift of Bor Kavčič). Resistance was flipped using pCP20 plasmid. *galK* under constitutive J23100 promoter was amplified from pKD13-PcgalK plasmid using primers 296 and 297 and transformed to replace the endogenous *fimH* gene using lambda red recombination. After recovery, any residual carbohydrate residues were removed by washing the cells several times with M9 buffer (Tomasek *et al,* 2018) before plating on M9 minimal medium containing 0.1 % galactose as only carbohydrate source for positive selection. Integration of *galK* gene into *fimH* was confirmed by PCR using primers 157 and 158. CFT073 *fimH* gene was amplified from a gblock (IDT) carrying the *fimH* sequence from CFT073 using primers 198 and 276 and integrated instead of the constitutive *galK* gene. After recovery, cells were washed several times as before. Transformants were counter-selected on artificial urine medium agar plates (AU-Siriraj; (Chutipongtanate & Thongboonkerd, 2010)) supplemented with 20 µg/ml L-aspartate and 20 µg/ml L-isoleucine (Bouvet *et al,* 2017), and containing 0.2 % 2-deoxy-D-galactose (DOG) and 0.2 % glycerol for the counter-selection. Pathogenic *fimH* integration was confirmed by PCR using primers 157 and 158. Resulting mutant was *E. coli W* locked-ON *fimH::fimHCFT073* – MG002.

##### FimH amino acid mutants

Single and triple point mutants of amino acids predicted to be most involved in binding to CD14 were generated using *galK* selection/counter-selection as mentioned above. Briefly, *galK* was deleted from CFT073 locked-ON mutants (KT179). Thereafter constitutive expressed *galK* was inserted in the endogenous *fimH* of this strain for selection. 100 µg gblocks (IDT) carrying either Y48A, R98A, T99A or the triple mutation (Y48A, R98A, T99A) in the *fimH* sequence from strain CFT073 were integrated instead of the constitutive *galK* gene. Correct integration of *fimH* having mutated amino acid residues was confirmed by sequencing. Resulting mutants were CFT073 locked-ON *fimHY48A R98A T99A* – KT260, CFT073 locked-ON *fimHY48A* – KT261, CFT073 locked-ON *fimHR98A* – KT262 and CFT073 locked-ON *fimHT99A* – KT263.

#### Inhibitors used

Since the suggested upper daily limit of orally applied D-mannose to treat UTIs is 9 g, leading to blood mannose levels of roughly 175 µM (Alton *et al,* 1997), we decided to compare this concentration to a strongly increased one of 30-60 g D-mannose resulting in 1 mM blood mannose levels. The small mannoside M4284 (medchemexpress) was used at 10 µM. The blocking CD14 antibody M14-23 was used at 20 µg/ml (Tsukamoto *et al,* 2010). Inhibitors were added as the same time as the bacteria, no pre-incubation steps were carried out.

### Method Details

#### Yeast agglutination assay

*Saccharomyces cerevisiae* was grown in YPD medium at 30 °C for 2 days. After centrifugation, cells were resuspended in M9 buffer to an OD_600_ of 1 and stored in the fridge. An aliquot of the yeast was transferred to glass slides and bacterial colonies were directly mixed into the yeast solution. Agglutination occurred within few seconds to 1 min. Pictures of agglutinated yeast and bacteria cells were taken on a brightfield microscope at 10x magnification and images were processed with Fiji.

#### Growth curve assay and doubling time estimation

Single bacterial colonies were inoculated in 160 µl R10H20 in 96-well plates and grown overnight at 220 rpm at 37 °C. The next day, cultures were diluted 1 in 1,000 in R10H20 supplemented with 0.0005 % triton-X and grown at 37 °C with shaking. Optical density was measured every 30 min at 600 nm at a Synergy H1 plate reader for a total of 7 h. The data were blank normalized, and the doubling time (dt) was calculated from exponential data using following formula 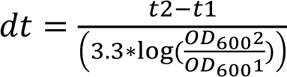.

For all assays, bacteria were grown to early-to mid-exponential phase (OD_600_ 0.25; *E. coli* W 4 h, CFT073 3 h 20 min; see Supplementary Figure 6A) in 1 ml R10H20 medium, if not indicated otherwise, at 37 °C in deep well plates.

#### Minimal inhibitory concentration (MIC) assay

Cultures were grown to OD_600_ of 0.25 and approximately 10^6^ bacteria were used for MIC assays. Serial dilutions of gentamicin were performed in a microdilution manner using 96-well plates and OD_600_ was measured after 18 h incubation at 37 °C with shaking. The threshold to calculate the MIC was set to detectable growth above the blank background after normalization. CFT073 locked-ON (KT179) and locked-OFF (KT180) mutants had similar MIC to gentamicin in R10H20 medium (see Supplementary Figure 6B). 5x the MIC was used to prevent extracellular growth of bacteria in the infection assays (7.5 µg/ml).

#### Electron microscopy

Bacteria were grown to mid-exponential phase in CAA M9 glycerol medium, fixed with glutaraldehyde (EM grade, final concentration 2.5 %) for 30 min at 4 °C, washed twice with PBS and concentrated in water. Formvar-coated copper grids were glow discharged for 2 min and 5*10^6^ bacteria were loaded for 5 min. Excess liquid was removed with filter papers and bacteria were stained with Uranyless for 2 min. After removal of excess liquid, the grids were washed 10 times in water and dried. EM images were taken at TEM T10 microscope at 80 kV. Images were processed with Fiji.

#### Predicted protein-protein interaction

Crystal structures of FimH (PDB: 6GTV), mouse CD14 (PDB: 1WWL), human CD14 (PDB: 4GLP), mouse TLR4 (PDB: 3VQ2) and mouse CD48 (PDB: 2PTV) were obtained from rscb.org, cleaned from solvents and other co-precipitated molecules using PyMol and run on HawkDock server to predict protein binding (Weng *et al,* 2019). Additional MM/GBSA analysis was run to predict free binding energy (Hou *et al,* 2011).

#### Bead binding assay

5 µl of 10 µm bead slurry (PureProteom Protein A magnetic beads, Merck) were used per reaction. Beads were washed 3x with PBS containing 0.005 % tween (PBS-T). Beads were collected using a magnetic stand. 5 µg CD14-Fc recombinant protein (RND System) in a total of 10 µl of PBS-T was coupled to the beads for 1 h at 4 °C with rotation. After washing the beads, 100 µl of bacteria grown to early-to mid-exponential phase in CAA M9 glycerol medium (∼1*10^7^ bacteria) were incubated in the presence of tween with the CD14-coupled beads for 1 h at 4 °C with rotation. After washing, this time with M9 glycerol medium containing tween, bead-bound bacteria were fixed with 0.5 % PFA in M9 buffer for 10 min at 4 °C.

For flow cytometry, samples were diluted in M9 buffer and analyzed on a FACS Canto II (BD). 10,000 events gated on FSC-A and SSC-A to exclude debris were recorded at medium flow rate and data were analyzed using FlowJo software. Three things should be noted: First, we observed that due to the high force applied in the sample injection tube during acquisition, the bacteria were separated from the magnetic beads they previously were bound to, leading to a single detectable fluorescent peak only. Second, due to their size, the magnetic beads are also detected in the gating range specific for the bacterial population (see Supplementary Figure 5C). Third, as can also be seen in the fluorescent images, the magnetic beads have weak auto-fluorescence in the FITC channel. We therefore quantified the amount of fluorescent events applying the same gating strategy as for the bacterial population only (Supplementary Figure 5B and C).

For fluorescent microscopy, samples were embedded in mounting buffer and spread on a cover slip. Images were taken with 100x magnification on a custom-built Olympus widefield microscope with Hamamatsu Orca Flash4.0v2 camera and LED-based fluorescence illuminator using YFP (x513/22,m543/22) fluorescence channel (Chait *et al,* 2017). Images were processed with Fiji and deconvoluted with Huygens software.

#### Type 1 pili extracts

Type 1 pili extracts were generated as described previously (Sheikh *et al,* 2017) with minor modifications. Briefly, CFT073 locked-ON (KT179) and CFT073 locked-ON *ΔfimH* (KT193) were grown overnight in CAA M9 glycerol medium and harvested at 4,000 g for 1 h. The cell pellet was resuspended in 1 mM Tris-HCl (pH 8.0) and incubated at 65 °C for 1 h with occasional vortexing. After pelleting the cells at 15,000 g for 10 min, type 1 pili were precipitated from the supernatant overnight in the presence of 300 mM NaCl and 100 mM MgCl_2_ at 4 °C. Type 1 pili were concentrated at 20,000 g for 10 min, washed once with 1 mM Tris-HCl and snap frozen in a small volume of 1 mM Tris-HCl.

##### Dot blot assay

The recombinant chimera CD14-Fc protein was biotinylated using the EZ-Link Sulfo NHS-LC-LC-Biotin kit (Thermofisher) and a 5-fold molar excess of biotin. In brief, CD14-Fc was dissolved in PBS at 1 mg/ml and biotinylated with a 5-fold molecular excess of biotin for 30 min at room temperature (RT). The biotinylation reaction was stopped with 3 mM Tris-HCl (pH 7.0).

The PVDF membrane was activated for 5 min with MeOH, washed for 5 min in water and allowed to dry for 5 min. 20 ng biotinylated CD14 and roughly 20 µg type 1 pili extract from CFT073 ON (KT179) and CFT073 ON *ΔfimH* (KT193) mutants were loaded onto the membrane. Protein spots were allowed to dry for 10 min and then the membrane was blocked in 3 % BSA in PBS for 30 min at 37 °C. 40 ng biotinylated CD14 in 3 % BSA in PBS and 0.05 % tween was blotted on the membrane and incubated for 1 h at 37 °C. Then the membrane was washed 3x in PBS with tween for 5 min each. Streptavidin-HRP antibody was pre-diluted 1:100 in 3 % BSA in PBS with tween and diluted once more 1:5,000 in PBS with tween. The membrane was incubated with Streptavidin-HRP for 1 h at RT. After washing again 3x as before, chemiluminescence was detected using clarity ECL substrates (Biorad).

#### Infection assays

DCs were seeded at a density of 1-2*10^5^ cells/ml in R10H20 medium (for adhesion assays black 24-well tissue-treated dishes were used, for any other assay non-treated dishes were used). DCs were matured either by addition of 200 ng/ml recombinant LPS or bacteria at a multiplicity of infection (MOI) of 10 (10 bacteria per 1 DC). 1 h post infection (pi) gentamicin was added at 7.5 µg/ml to prevent extracellular growth of bacteria. 18-20 h pi subsequent assays were performed.

#### Adhesion assay

Non-adherent DCs were removed, and adherent cells were washed twice with 500 µl PBS. Adherent cells were stained with Hoechst 33342 (NucBlue reagent, 2 drops/ml) in R10H20 medium for 30 min at 37 °C. Cells were washed twice and 1 ml Live Cell Imaging solution (140 mM NaCl, 2.5 mM KCl, 1.8 mM CaCl_2_, 1.0 mM MgCl_2_, 20 mM HEPES, pH 7.4) was added to the wells. Fluorescence was measured with a Synergy H1 plate reader (excitation 490 nm, emission 520 nm, bottom reading without lid, 50 data points per well). Pictures of adherent cells were taken on a brightfield microscope at 4x magnification and images were processed with Fiji.

#### Flow cytometry staining

DCs were collected and incubated in FACS buffer (1x PBS, 2 mM EDTA, 1 % BSA; RT) or Tyrodés buffer (used for active CD11b and β1 staining, on ice) with Fc receptor block for 20 min. Cells were stained for 30 min with antibodies using the respective buffer with Fc receptor block. Cells were washed twice with PBS and resuspended in the respective buffer for analysis on FACS Canto II (BD). 10,000 events gated on FSC-A and SSC-A to exclude debris were recorded at medium flow rate. Data were analyzed using FlowJo software by performing doublet discrimination. Antibodies used are listed in Table 3.

#### Ex vivo ear crawl out assays

Ear crawl out assays were performed similar as published previously with minor modifications (Kopf *et al,* 2020). In brief, ears from 5-week-old female C57Bl/6J WT mice were first UV sterilized for 10 min and then split into dorsal and ventral halves. Ventral halves were placed in R10H20 medium, ventricles facing down. Ears were incubated with 10^6^ CFT073 locked-ON (KT179) or OFF (KT180) bacteria for 48 h, renewing the infection stimulus after 24 h. 1 h after every infection 7.5 µg/ml gentamicin was added to the medium. Ears were fixed using 4 % PFA and immersed using 0.2 % triton-X. After blocking in 1 % BSA in PBS, lymphatics were stained for 90 min using rat anti-Lyve-1 antibody and DCs were stained using biotinylated anti-MHCII antibody. Secondary antibodies, anti-rat F(ab’)2-AF488 and Streptavidin-AF647, were used subsequently for 45 min each. Ears were fixed on cover slips with ventricles facing up using cover glasses. 10 µm z-stacks were taken on inverted LSM800 confocal microscope with 488 and 640 nm LED-laser light source. Images were taken from 3 biological replicates (except *tlr4*^-/-^ where 2 biological replicates were imaged) analyzing at least 2 field of views each. Maximum intensity projection images were processed with Fiji. Images were analyzed using custom-made scripts in Fiji. Pre-processing was done using lymphatics script and analysis using LVmeanDCarea script.

#### In vitro 3D collagen migration assay

3D collagen chemotaxis assays were performed as described previously (Leithner *et al,* 2016), with minor modifications. Assays were performed in PureCol bovine collagen with a final collagen concentration of 1.6 mg/ml in 1x minimum essential medium eagle (MEM) and 0.4 % sodium bicarbonate using 1-2*10^5^ DCs. The collagen-cell mixture was cast to custom-made migration chambers and polymerized for 1 h at 37 °C. CCL19 chemokine (RND Systems) in R10 (0.625 µg/ml) was pipetted on top of the gel and the chambers were sealed with paraffin. Images were taken every 30 sec for a total of 5 h on bright field microscopes using 4x magnification and an exposure of 20 ms. Data were analyzes using custom-made Fiji scripts: images were pre-processed using Tracking_pre-processing_for_brightfield script and analyzed using migrationspeedREP script.

#### In vitro extracellular matrix migration assay

Cell-derived matrixes (CDM) were produced as described previously (Kaukonen *et al,* 2017). In brief, round shaped coverslips were coated with 0.2 % gelatin in PBS in 24-well dishes for 1 h at 37 °C. Gelatin was crosslinked with 1 % glutaraldehyde in PBS for 30 min at RT and quenched with 1 M glycine in PBS for 20 min at RT. After washing the coverslips twice with PBS, 5*10^4^ 3T3 mouse fibroblasts in DMEM, GlutaMAX, supplemented with 10 % FCS, 100 U/ml penicillin and 100 µg/ml streptomycin were seeded per well. After 48 h 3T3 fibroblasts reached confluency and were treated daily with ascorbic acid for better crosslinking of the extracellular matrix. Old medium was gently removed and fresh medium with 50 µg/ml sterile ascorbic acid was added for 10-14 days. 3T3 fibroblasts were extracted with extraction buffer (0.5 % Triton-X, 20 mM NH4OH in PBS) for 2 min and washed twice with PBS containing 1 mM CaCl_2_ and 1 mM MgCl_2_ (PBS/Ca/Mg). DNA was digested with 100 µg/ml DNaseI in PBS/Ca/Mg for 1 h at 37 °C and CDM were washed twice with PBS/Ca/Mg before storage in PBS/Ca/Mg supplemented with 100 U/ml penicillin and 100 µg/ml streptomycin at 4 °C.

Before use, CDM were placed onto custom-made imaging chambers and incubated with R10 medium for 1 h at 37 °C. The medium was removed and a 1 µl CCl_2_1 chemokine (RND Systems; 25 µg/ml) spot was injected into the CDM and incubated for 10 min. 1 ml R10 medium was added on top and the CDM was incubated for 1 h at 37 °C to allow a chemokine gradient to form. After washing twice gently with R10 medium, CFT073 locked-ON (KT179) and locked-OFF (KT180) stimulated DCs were concentrated by centrifugation and the dense cell pellet was pipetted into the CDM at the opposite site to the chemokine spot. 2 ml of R10 medium was added and images were taken every minute for a total of 6 h on a bright field microscope using 10x magnification and an exposure of 20 ms. Single cells outside of the cell cluster were counted after 5 h. Images were processed with Fiji.

#### In vitro T cell assay

T cell assays were performed as described previously (Leithner *et al,* 2021). In brief, primary naïve CD4^+^ T cells were isolated from the spleen of OT-II mice (B6.Cg-Tg(TcraTcrb)425Cbn/J) using EasySep Mouse CD4^+^ T cell isolation kit (Stemcell Technologies) after homogenization with a 70 µm cell strainer and resuspending the cells in PBS supplemented with 2 % FCS and 1 mM EDTA. T cells were co-cultured with DCs matured with CFT073 locked-ON (KT179) or OFF (KT180) at a ratio of 5:1 (5*10^4^ T cells:1*10^4^ DCs) in 96-well round bottom well plates in R10 medium.

#### T cell activation

After 24 h co-culture in the presence of 0.1 µg/ml ovalbumin (OVA), medium was removed by spinning. Cells were incubated with Fc receptor block in FACS buffer and stained with anti-CD4, anti-CD69 and anti-CD62-L antibodies for 15 min at 4 °C. After resuspending cells in FACS buffer, 100 µl were recorded on FACS Canto II (BD) and the ratio of CD69 to CD62L expression of CD4^+^ T cells was analyzed by FlowJo software.

#### T cell priming

T cells were stained with 5 µM CFSE stain in 5 % FCS in PBS for 5 min at RT. After 30 min recovery in R10 medium at 37 °C, cells were routinely checked for fluorescence.

After co-culturing with DCs for 4 days in the presence of 0.1 µg/ml OVA, medium was removed by spinning. Cells were stained with anti-CD4 antibody for 10 min at 4 °C. After resuspending cells in FACS buffer and 7AAD viability stain, 100 µl were recorded on FACS Canto II (BD). Dividing T cells were analyzed with FlowJo software.

#### DC-T cell interaction time

Interaction time of DCs and T cells was measured as described previously (Leithner *et al,* 2021). In brief, glass bottom dishes were plasma cleaned for 2 min and coated with 1x poly-L-lysine in water for 10 min at RT. Dishes were washed two times with water and dried overnight. 1.5*10^5^ DCs, pre-loaded with 0.1 µg/ml OVA for 1 h 30 min, were mixed with 3*10^5^ T cells and loaded onto the coated dishes in a total volume of 300 µl. Images were taken every 30 sec for a total of 6 h on bright field microscope using 20x magnification and an exposure of 20 ms. Images were processed with Fiji.

#### Quantification and Statistical Analysis

Data are represented as means ± standard deviations. Statistics were performed using GraphPad Prism version 9.0.2 for Windows. Statistical details for each experiment can be found in the respective Figure legends. Significance was defined as follows: * p<0.1, ** p<0.05, *** p<0.01, **** p<0.001.

## Acknowledgements

We thank Ulrich Dobrindt for providing UPEC strain CFT073, Vlad Gavra and Maximilian Götz, Bor Kavčič, Jonna Alanko and Eva Kiermaier for help with experiments and Robert Hauschild, Julian Stopp and Saren Tasciyan for help with data analysis. We thank the IST Austria Scientific Service Units, especially the Bioimaging facility, the Preclinical facility and the Electron microscopy facility for technical support, Jakob Wallner and all members of the Guet and Sixt lab for fruitful discussions and Daria Siekhaus for critically reading the manuscript. This work was supported by grants from the Austrian Research Promotion Agency (FEMtech 868984) to I.G., the European Research Council (CoG 724373) and the Austrian Science Fund (FWF P29911) to M.S.

## Author Contributions

Conceptualization, K.T., A.L., M.S.L., C.C.G. and M.S.; Methodology, K.T., C.C.G. and M.S.; Validation, K.T., C.C.G. and M.S.; Investigation, K.T., A.L., I.G.; Formal analysis, K.T.; Visualization, K.T.; Supervision, C.C.G. and M.S.; Writing – original draft, K.T.; Writing – review & editing, K.T., C.C.G. and M.S. with the help of all authors; Funding acquisition – C.C.G. and M.S.

## Data and material availability

Further information and requests for resources, scripts and data should be directed to Călin Guet and Michael Sixt.

## Supplemental Items

**Table S1.** Predicted free binding energies of mouse CD14, TLR4 and CD48 to FimH.

**Table S2.** Amino acids of FimH responsible for binding mouse CD14 and mannose, and amino acids of mouse CD14 responsible for FimH and LPS binding.

**Table S3.** Predicted free binding energy of human CD14 and FimH.

**File S1.** PDB file of mouse CD14 and FimH.

**File S2.** PDB file of human CD14 and FimH.

